# A longitudinal analysis of puberty-related cortical development

**DOI:** 10.1101/2020.11.05.370460

**Authors:** Nandita Vijayakumar, George Youssef, Nicholas B Allen, Vicki Anderson, Daryl Efron, Philip Hazell, Lisa Mundy, Jan M Nicholson, George Patton, Marc L Seal, Julian G Simmons, Sarah Whittle, Tim Silk

**Author notes:** Co-senior authors (Sarah Whittle, Tim Silk). Corresponding author: Nandita Vijayakumar, School of Psychology, Deakin University, 221 Burwood Highway, Burwood, VIC 3125.

## Abstract

The brain undergoes extensive structural changes during adolescence, concurrent to puberty-related physical and hormonal changes. While animal research suggests these biological processes are related to one another, our knowledge of brain development in humans is largely based on age-related processes. Thus, the current study characterized puberty-related changes in human brain structure, by combining data from two longitudinal neuroimaging cohorts. Beyond normative changes in cortical thickness, we examined whether individual differences in the rate of pubertal maturation (or “pubertal tempo”) was associated with variations in cortical trajectories. Participants (N = 192; scans = 366) completed up to three waves of MRI assessments between 8.5 and 14.5 years of age, as well as questionnaire assessments of pubertal stage at each wave. Generalized additive mixture models were used to characterize trajectories of cortical development. Results revealed widespread linear puberty-related changes across much of the cortex. Many of these changes, particularly within the frontal and parietal cortices, were independent of age-related development. Males exhibiting faster pubertal tempo demonstrated greater thinning in the precuneus and frontal cortices than same-aged and -sex peers. Findings suggest that the unique influence of puberty on cortical development may be more extensive than previously identified, and also emphasize important individual differences in the coupling of these developmental processes.

---

The transition from childhood to adolescence marks a period of prolonged cortical thinning that extends into the mid 20s (Mills et al., 2016; Tamnes et al., 2017). While most of this literature has focused on age-related changes in brain structure during adolescence, this period is also defined by pubertal development. The release of pubertal hormones trigger the process of sexual maturation, and it has long been theorized that these hormonal changes also play a role in determining adolescents’ emotions and behaviours via their actions on the developing brain (Blakemore et al., 2010; Dahl, 2004). Importantly, individual differences in pubertal development are better predictors of many socioemotional outcomes than age (Ullsperger & Nikolas, 2017), reflecting the importance of considering puberty-related changes in underlying neural circuits. Animal research and a growing body of work in humans shows a link between pubertal development and cortical maturation (Herting & Sowell, 2017; Juraska & Willing, 2017; Vijayakumar et al., 2018). However, there is a need for longitudinal research in humans to examine the coupling of developmental trends in these biological processes, which may be obscured in cross-sectional studies. Importantly, such a design is best suited to distinguish the potential effects of pubertal stage and chronological age on brain development.

Pubertal development occurs in two phases, adrenarche and gonadarche. Adrenarche, the earlier phase of puberty, begins when changes in the hypothalamic-pituitary-adrenal axis produce androgens that are responsible for secondary sex characteristics, such as pubic hair growth, body odor and acne (Havelock et al., 2004). Gonadarche is triggered by the activation of the hypothalamic-pituitary-gonadal axis; the hypothalamus releases substantial amounts of gonadotropin-releasing hormone that triggers the pituitary to produce follicle stimulating and luteinizing hormones, which in turn stimulate the ovaries and testes to produce sex steroid hormones that are responsible for reproductive maturity and other secondary sex characteristics (Plant & Barker-Gibb, 2004; Veldhuis, 1996). These sex hormones also act via androgen and estrogen receptors in the brain to modulate the synthesis, release and metabolism of various neurotransmitters (e.g., noradrenaline, dopamine, serotonin, glutamate, and GABA) and associated neuropeptides that influence the excitability, synaptic function, and morphology of neurons (Juraska & Willing, 2017). Thus, there is growing support for being puberty a period when hormones exert “re-organizational” effects on brain structure (Schulz et al., 2009).

Although puberty comprises a multitude of biological and physical changes, the most common conceptualization of puberty is the progression through the five “Tanner” stages of physical development based on changes in the breast (for females), genitalia, and pubic hair (Tanner, 1962). Prior neuroimaging research indicates a general pattern of reductions in grey matter thickness within prefrontal and parietal cortices (KoolschiJn et al., 2014; Peper et al., 2009; Pfefferbaum et al., 2015), and to a lesser extent changes in parts of the medial temporal lobe (Hu et al., 2013), as individuals progress through pubertal stages. However, these studies have used cross-sectional samples, thus limiting conclusions about developmental processes (as discussed in Herting & Sowell, 2017; Vijayakumar et al., 2018). Additionally, they may be under-powered to identify unique pubertal effects, as controlling for age can remove much of the between person variance in cross-sectional designs (due to the strong correlation between age and pubertal stage). There have been a few longitudinal studies examining the relationship between puberty and subcortical development, which have identified changes in subcortical volumes with pubertal stage, as well as qualitative and quantitative sex differences (Goddings et al., 2014; Herting et al., 2014; Wierenga et al., 2018). Importantly, pubertal stage has been found to be a better predictor of development of a number of subcortical structures than age (Wierenga et al., 2018), and sex hormones (i.e., testosterone, estradiol) have been implicated (Herting et al., 2014; Wierenga et al., 2018). However, longitudinal examination of puberty-related cortical development has been limited to a single sample of 281 children and adolescents (with 469 scans between 4 and 22 years of age). Using this sample, Nguyen and colleagues (2013) identified significant negative associations between testosterone and cortical thickness in more developed adolescents (pubertal stages 3-5), including the left posterior cingulate, precuneus, dorsolateral prefrontal, and anterior cingulate cortices in males, and right somatosensory cortex in females. They also found stronger negative associations between pubertal stage and cortical thickness in the later stages (i.e., 3 to 5) relative to earlier stages (i.e., 1 to 2), but did not report whether specific cortical regions were involved. Thus, further longitudinal research is needed to improve our understanding of the role of puberty in cortical development.

Aside from pubertal stage, inter-individual differences in pubertal maturation can be described using the concepts of pubertal timing and tempo. Timing refers to the stage of an individual relative to same-sex and -age peers, while tempo refers to the rate of progression through the pubertal stages relative to same sex- and age-matched peers (Marceau et al., 2011). Earlier timing and faster tempo have been found to predict internalizing and externalizing symptoms in both sexes (Beltz et al., 2014; Marceau et al., 2011; Mendle et al., 2010), and thus consideration of inter-individual differences in pubertal maturation may provide insight into the neurodevelopmental correlates of socioemotional behaviors and psychopathology. A few studies have investigated the neural correlates of pubertal timing, identifying decreased frontal and temporal thickness with earlier timing (Hu et al., 2013; Peper et al., 2009). However, given that longitudinal designs are necessary to study pubertal tempo, to date, there has been little consideration of how it may relate to variation in cortical developmental trajectories. Herting and colleagues (2015) present the only study to do so.

Using a sample of 81 adolescents with 162 scans between 10 – 16 years, they found that greater increase in pubertal stage over 2 years (i.e., faster tempo) was related to less thinning of the superior frontal and right superior temporal gyri, with the latter effect being stronger in females. Further consideration of such variability is necessary to improve our understanding of deviations from normative development, including whether pubertal tempo represents a potential biological indicator of risk for behavioral and emotional problems during adolescence.

The current study addresses limitations in the pubertal neuroimaging literature using data from two longitudinal community-based cohort studies, which assessed children up to three times between late childhood and mid-adolescence (8.5 to 14.5 years of age). We examined changes in cortical thickness with pubertal maturation. Based on prior cross-sectional and longitudinal literature, we hypothesized that there would be reductions in cortical thickness with increasing pubertal stage, particularly within prefrontal and parietal cortices. We also conducted exploratory examination of sex differences in these trajectories, but did not have specific hypotheses regarding such effects. Finally, we examined whether pubertal tempo (i.e., rate of pubertal maturation) was related to cortical developmental trajectories, hypothesizing that faster tempo would be related to lesser reductions in cortical thickness based on preliminary findings in the literature. Note that the current dataset was not suited to examine pubertal timing as discussed in the limitations section.

## Methods

### Participants

Participants were based in Melbourne, Australia, and were recruited into one of two longitudinal cohorts: i) Neuroimaging of the Children’s Attention Project (NICAP), and ii) imaging brain development in the Childhood to Adolescence Transition Study (iCATS). NICAP participants were recruited as typically developing controls into a community-based study of children with and without ADHD. iCATS participants were recruited based on adrenal hormone levels (dehydroepiandrosterone and testosterone), with the intent to maximize variance in hormone levels and pubertal development in the sample. Further details on the NICAP and iCATS cohorts are presented in Silk et al., (2016) and Simmons et al., (2014), respectively. Exclusion criteria for these analyses included MRI contraindications, developmental disability, history of a neurological or serious medical disorder (e.g., diabetes, kidney disease), and concurrent use of psychotropic medications. For both cohorts, written informed consent was obtained from the parent/guardian of all participants, and ethics approval was granted by The Royal Children’s Hospital Human Research Ethics Committee, Melbourne (NICAP #34071; iCATS #32171). Protocols were also ratified by the Human Research Ethics Committees of Deakin University (NICAP #2016-394) and The University of Melbourne (iCATS #1238745).

The NICAP cohort underwent up to 3 assessments between the ages of 9.5 and 14.5 years, with two approximately 18-month intervals (M = 1.432, SD = 0.222, 1.021 - 2.330 years) between assessments. The iCATS cohort underwent 2 repeated assessments between the ages of 8.5 and 13.5 years, with one approximately 36-month interval (M = 2.763, SD = 0.243, 2.158 - 3.344 years) between assessments. At baseline (wave 1) the two cohorts did not differ in biological sex (χ^2^ = 1.342, df = 1, p = 0.247), pubertal stage (Mean: NICAP = 1.316, iCATS = 1.282, t_(161)_ = −0.431, p = 0.666), or intelligence (based on Wechsler Abbreviated Scale of Intelligence – Matrix Reasoning T-score; Mean: NICAP = 52.607, iCATS = 54.096, t_(179)_ = 1.225, p = 0.222). However, the iCATS sample was significantly younger than the NICAP sample at baseline (Mean: NICAP = 10.425, iCATS = 9.556, t_(157)_ = −14.928, p < 0.001) and had higher socioeconomic status (SES; based on Socio-Economic Indexes for Areas - Index of Relative Socio-economic Advantage and Disadvantage [based on Australian Census data]; Mean: NICAP = 1018.326, iCATS = 1056.175, t_(198)_ = 4.887, p < 0.001). In total, 192 participants (96 males, 90 NICAP) were included, with a total of 366 scans (186 males, 207 NICAP). Note that both cohorts only examined biological sex (not gender identity). A visual representation of the age and Tanner stage at each assessment point for each participant in this combined sample is presented in Figure 1 (refer to Figures S1 and S2 in the Supplementary materials for a breakdown by cohort).

**Figure 1.**
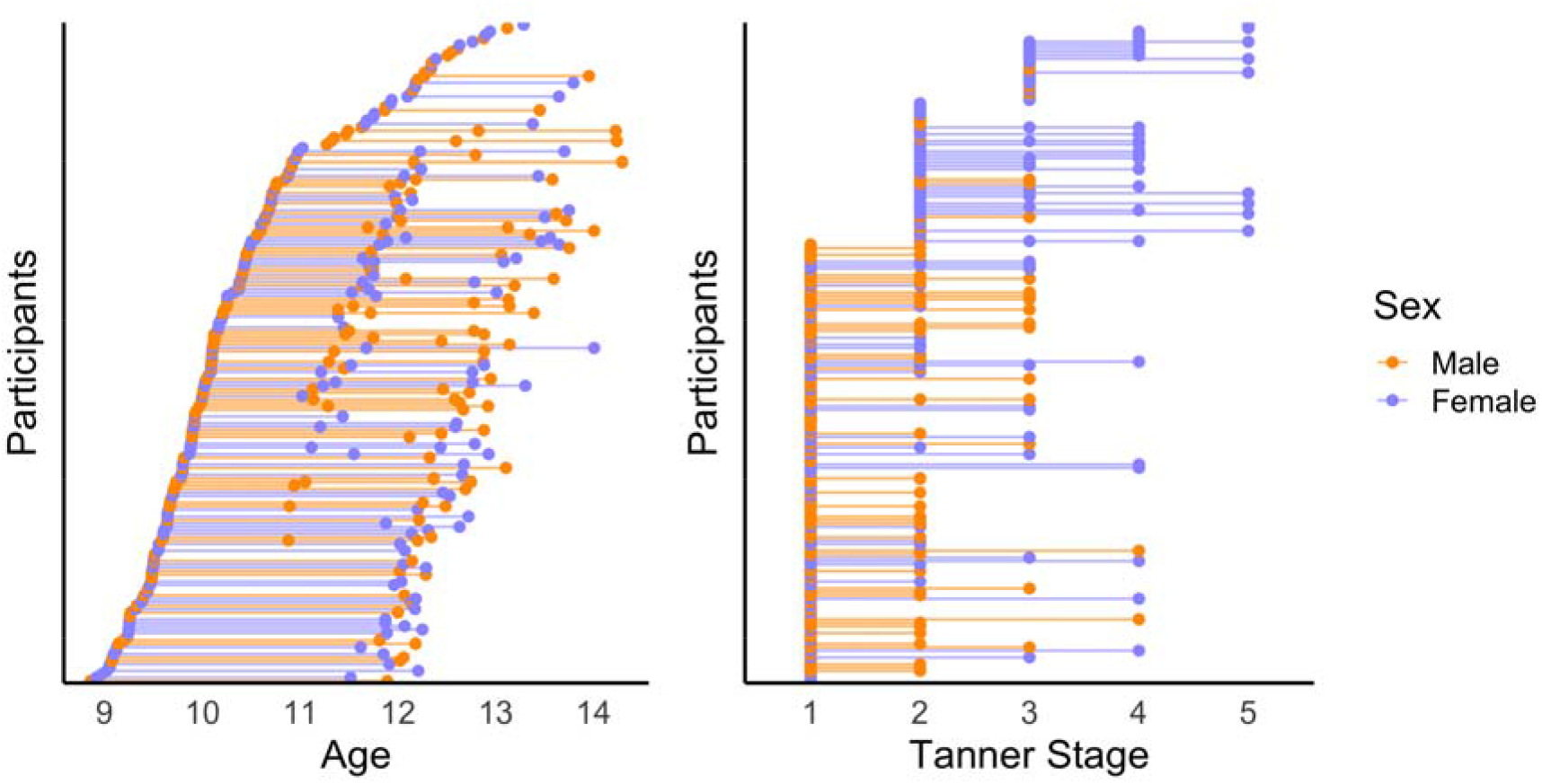
Distribution of sample by age and pubertal (Tanner) stage.

### MRI Acquisition & Processing

In both cohorts, participants were given information on MRI (including a video) prior to their assessment, in order to familiarize them with the procedure and minimize anxiety as much as possible. Participants completed a mock MRI scan before their actual scan at wave 1 (repeated at subsequent waves if the participant wished or the researcher deemed it appropriate). Neuroimaging data for both cohorts were acquired at a single site, on a 3 Tesla Siemens scanner (Siemens, Erlangen, Germany) at the Murdoch Children’s Research Institute in Melbourne, Australia. Both waves of iCATS, and waves 1 and 2 of NICAP, were collected on a TIM Trio scanner. The final wave of NICAP was collected after an upgrade to a MAGNETOM Prisma scanner, which has been accounted for within statistical modelling. Refer to the Supplement for analyses investigating the potential effects of scanner upgrade.

Participants lay supine in a 32-channel head coil during the MRI scan. Structural T1-weighted images were acquired as follows: *NICAP* – MEMPRAGE with repetition time = 2530 ms, echo time = 1.77, 3.51, 5.32, 7.2ms, flip angle = 7 °, field of view = 230 mm^2^, resulting in 176 contiguous slices with voxel dimensions 0.9 mm^3^; *iCATS* – MPRAGE with repetition time = 1900 ms, echo time = 2.24 ms, flip angle = 9 °, field of view = 230 mm^2^, resulting in 176 contiguous slices with voxel dimensions 0.9 mm^3^.

T1-weighted images were processed through FreeSurfer 6.0, a freely available image analysis suite for cortical reconstruction and volumetric segmentation (http://surfer.nmr.mgh.harvard.edu/). Specifically, images were processed with the submillimeter reconstruction (Zaretskaya et al., 2018) and the longitudinal stream that creates an unbiased within-subject template space from all available data using robust, inverse consistent registration. The template is used as an estimate to initialize segmentation processes for each time point, providing common information regarding anatomical structures, and has been found to significantly increase reliability and statistical power (Reuter et al., 2012; Reuter & Fischl, 2011). The quality of i) raw images and ii) (longitudinal) cortical reconstructions was visually inspected and rated for all scans. Raw images were rated on a 4-point scale, with ratings of “3” and “4” excluded. Processed images were rated on a 3-point scale, with ratings of “3” excluded. Images were also processed through MRIQC (v0.14.2) to supplement the visual inspection (Esteban et al., 2017). No manual edits were made to the remaining (included) data. Refer to the Supplement for further detail on quality control procedures, including the visual rating process. Mean cortical thickness estimates from 62 regions of the Desikan-Killiany-Tourville (DKT) atlas were extracted and used in subsequent analyses.

### Puberty

Pubertal stage was measured at each time point (across both cohorts) using the parent-report Sexual Maturity Status (Morris & Udry, 1980). Recognizing that physical examination is not necessarily a gold standard measure of puberty, correlations between parent report of Tanner stage and physical exam range from 0.75 to 0.87, suggesting good validity (Chavarro et al., 2017). Further, correlations are stronger for children/adolescents at lower Tanner stages (Dorn et al., 1990), suggesting that validity may be higher in the current sample. The survey consists of a series of stylized line drawings of girls/boys bodies at differing stages of pubertal development. Parents with a female child looked at a page with five stages of breast development, and five stages of hip and pubic hair development, and circled a number above the two series of images that most accurately represented their daughter’s development. Parents with a male child completed the same task, but with a single series of images of the five stages of male genital and pubic hair development. These images directly correspond to the five Tanner stages of pubertal development and have shown to have good reliability with physician ratings (Dorn & Biro, 2011). For females, the higher score for breast or pubic hair development images was used if there was a discrepancy (N = 69, 38%). Additionally, for females that only had data for either breast or pubic hair (N = 9, 5%), the single available score was used to measure Tanner stage.

### Statistical analyses

#### Cortical development

First, multiple imputation was undertaken using “mice” (Buuren & Groothuis-Oudshoorn, 2011) in R (R Core Team, 2013) to deal with missing Tanner stage. There were 21 data points missing in NICAP (12%, 15 males): 5 at wave 1, 5 at wave 2, and 10 at wave 3. There were 7 data points missing in iCATS (4%, 5 males): 4 at wave 1 and 3 at wave 2. Imputation was conducted on a long-format data set, using a proportional odds model for the ordinal Tanner stage variable. A total of 30 imputations were run, and the average (mode) value across these imputations was used as the final Tanner stage. This was done separately for NICAP and iCATS as different predictor variables were available to perform imputation in each cohort. The imputation model for NICAP included sex, age and the Pubertal Development Scale (Petersen et al., 1988), while iCATS included sex, age and pubertal hormone levels (specifically testosterone, dehydroepiandrosterone (DHEA) and DHEA-sulphate). Refer to the supplement for further detail on these additional pubertal measures that were used during imputation.

Next, generalized additive mixed models were used to examine the development of cortical thickness in relation to Tanner stage (i.e., TS). As an exploratory aim of this study, we also investigated sex differences in puberty-related cortical development. To do so, the following models were examined using the “gam” function within the “mgcv” package (Wood, 2006) in R:

1. *Null:* thickness ~ cohort + scanner + sex + s(id, bs = “re”)
2. *Puberty-related nonlinear cortical development:* thickness ~ cohort + scanner + sex + s(id, bs = “re”) + s(TS, bs = “cs”, k = 3)
3. *Puberty-related linear cortical development:* thickness ~ cohort + scanner + sex + s(id, bs = “re”) + TS
4. *Sex differences in puberty-related nonlinear cortical development:* thickness ~ cohort + scanner + sex + s(id, bs = “re”) + s(TS, bs = “cs”, k = 3) + s(TS, by = sex, bs = “cs”, k = 3)
5. *Sex differences in puberty-related linear cortical development:* thickness ~ cohort + scanner + sex + s(id, bs = “re”) + TS + TS*sex

The “s(id, bs = “re”)” term represents the random intercept for each individual, while the s(TS, bs = “cs”, k = 3) term represents developmental *smooth* terms using a penalized cubic regression spline and a basis function of 3 (chosen given the constrained age span that was investigated). Cohort (iCATS, NICAP), scanner (pre-, post-upgrade), and sex (males, females) were included as binary factors. Sex was also an ordered factor, in order to examine the developmental (trajectory) difference between males and females in Model 4. All models were examined with maximum likelihood (ML) estimation. This model building procedure was repeated for each of the 62 regions of the DKT atlas.

A series of model comparisons was undertaken to determine the most parsimonious model fit across Models 1-5. To do so, we used the compareML function within “itsadug” (van Rij et al., 2017), which compares two models on the basis of the minimized smoothing parameter selection score (i.e., “ML”), with a χ^2^ test on the difference in scores and degrees of freedom (that provided a p-value indicating whether the *more complex model* provided better fit to the data than the less complex model). First, we examined developmental trajectories across the group (i.e., males and females) by comparing Models 1 and 2 (i.e., null vs. puberty-related nonlinear cortical development). Additionally, we checked whether the *smooth* term for Tanner stage in Model 2 was significant following correction for multiple comparisons using False Discovery Rate 0.05, to account for the 62 regions that were examined. Next, we examined the significance of nonlinearity in cortical trajectories by comparing Models 2 and 3 (i.e., puberty-related *non-linear vs. linear* cortical development). This was undertaken as the “gam” function with a penalized cubic regression spline does not provide a statistical differentiation between linear and nonlinear trends (i.e., plotting the predicted values from gam may suggest a linear trend, but it is uncertain whether a linear or non-linear trend better provides a better fit).

For exploratory analyses of sex differences, we compared Models 2 and 4 to determine whether males and females differed in puberty-related cortical trajectories. We chose to compare Models 2 with 4 even for regions that showed linear trends across both sexes as the “gam” function will fit a linear *trend* when it is the best fit. Next, we statistically examined the significance of nonlinear trends in sex differences by comparing Models 4 and 5 (i.e., sex differences in puberty-related *nonlinear vs. linear* cortical development).

Finally, we examined puberty-related cortical development independent of age-related development by re-running Models 1 to 4 including a *smooth* term of age (i.e., s(age, bs = “cs”, k = 3)).

#### Pubertal tempo

Based on prior literature examining pubertal tempo using growth models (Marceau et al., 2011; Mendle et al., 2010), we calculated pubertal tempo for each individual using linear mixed models (in Stata 15; StataCorp, 2017). These predicted Tanner stage from age and age^2^, with random effect estimates for id and age (i.e., TS ~ age + age^2^, random effect = [1 + age | id]). Models were estimated for each sex separately. Random age slopes were extracted for each individual as an index of pubertal tempo as they reflect individual differences in the linear age-slope relative to the group. Thus, higher random slope values suggest that individuals were progressing through the Tanner stages faster than their peers over time (i.e., steeper positive trajectory than the group-level “fixed-effect” trajectory), while lower random slopes suggest that individuals were slower at progressing through the Tanner stages relative to their peers (i.e., shallower positive trajectory relative to the group-level trajectory). These random slopes are estimated from all available data, and thus account for differences in time intervals across participants. Note that these analyses were conducted on non-imputed data, given that age was itself a predictor in the imputation models described above. Random slopes were available for all but 1 female and 1 male; a total of 95 males (185 scans) and 95 females (178 scans). This sample did not differ from the full sample (N = 192) in age (t(1.01) = 1.61, p = 0.35), sex (X^2^ (1) = 0, p = 1), or cohort (X^2^ (1) = 0.65, p = 0.42) distributions.

Next, the random slopes were used to examine the development of cortical thickness in relation to pubertal tempo. These analyses were conducted within each sex given that random slopes in males and females were conditional on separate models, and therefore absolute values were not comparable across sexes. Within each sex, we examined whether changes in cortical thickness over time differed as a function of pubertal tempo using the following model:

6) *Pubertal tempo-related variability in cortical development:* thickness ~ cohort + scanner + tempo + s(age, bs = “cs”, k = 3) + s(age, by = tempo, bs = “cs”, k = 3) + s(id, bs = “re”)

The s(age, by = tempo, bs = “cs”, k = 3) term in Model 6 represents a linear interaction between the *smooth* age term and tempo, and informs us whether individual differences in pubertal tempo are associated with different rates of cortical development over age/time. CompareML was used to determine whether the interaction term (i.e., Model 6) improved model fit beyond the main effect of tempo (Model 7: thickness ~ cohort + scanner + tempo + s(age, bs = “cs”, k = 3) + s(id, bs = “re”)) and a null model without tempo (Model 8: thickness ~ cohort + scanner + s(age, bs = “cs”, k = 3) + s(id, bs = “re”)). Additionally, we checked whether the s(age, by = tempo, bs = “cs”, k = 3) term in Model 6 was significant following correction for multiple comparisons using False Discovery Rate 0.05, to account for the 62 regions that were examined.

Models for females also controlled for individual differences in Tanner stage at baseline (i.e., random intercept), as it was significantly positively correlated to random slopes (see Results section for further details). We specifically incorporated a main effect of pubertal stage at baseline and its interaction with age (i.e., “random intercept + s(age, by = random intercept, bs = “cs”, k = 3)”) in the null, main effect, and intercept models. As there was almost no variance in random intercept in males (SD < 0.001), we did not incorporate this variable into models for males.

#### Sensitivity analyses

A series of sensitivity analyses were conducted. First, we included body-mass index (BMI) and SES as additional covariates of non-interest to models. Second, we re-ran analyses with the exclusion of 11 participants on steroid medications (males = 6, NICAP = 0). Third, we reran analyses with the exclusion of 5 participants who exhibited reductions in Tanner stage over time (males = 5, NICAP = 2).

## Results

### Cortical development and pubertal stage

Model comparisons revealed puberty-related trajectories across most of the cortex, with the only exceptions being the bilateral entorhinal cortices. Findings were consistent when considering the significance of the *smooth* term for Tanner stage following FDR correction. Trajectories across the cortical mantle were better characterised by linear than nonlinear reductions. On average, cortical regions exhibited 1.29% reduction (SD: 0.43; range: −2.00 to 0.04) in thickness per Tanner stage (see Figure 3). Refer to Table S6 for model comparisons, and Table S7 for summary statistics of the linear model. An illustration of effect sizes by sex are presented in Figure S8.

**Figure 3.**
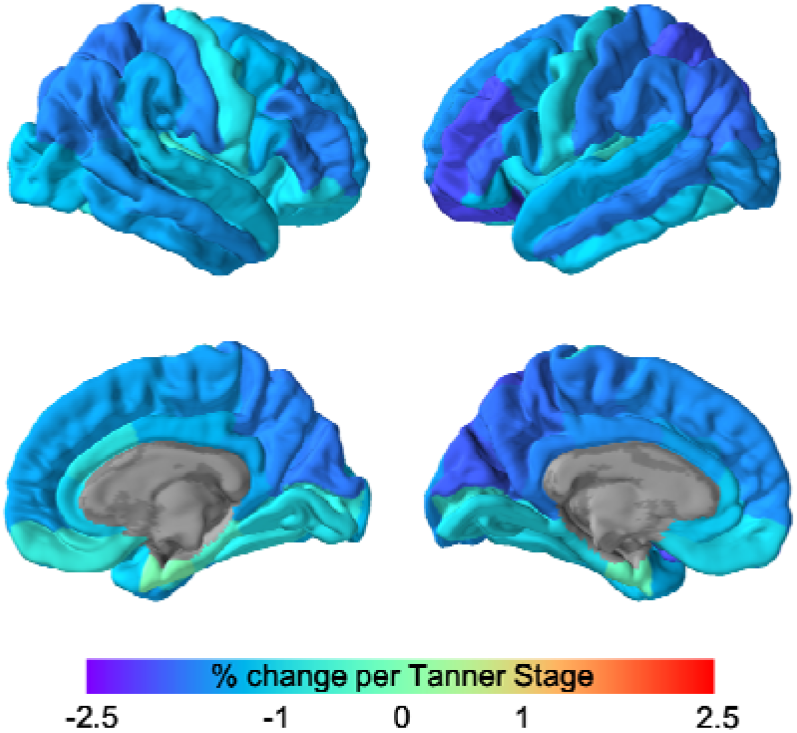
Puberty-related changes in cortical thickness, without controlling for age. Values represent percent change per Tanner stage (relative to the average thickness at wave 1).

Analyses revealed significant linear reductions in thickness with age across most of the cortical mantle (refer to the Supplement for further detail, with results presented in Tables S8 and S9). Model comparisons also revealed that a number of regions exhibited puberty-related cortical changes over and above these age-related changes (i.e., when controlling for neurodevelopmental variance associated with age; see Figure 4a). This included much of the bilateral frontal cortex extending medially to the mid and (left) posterior cingulate and precuneus, as well as extending laterally to the parietal and parts of the temporal cortices (these regions also exhibited significant *smooth* terms for Tanner stage following FDR correction for multiple comparisons). Conversely, regions that did not exhibit puberty-related development when controlling for age clustered primarily within parts of the anterior cingulate, occipital and temporal cortices. Again, model comparisons revealed that significant puberty-related trajectories were best characterised by linear reductions in thickness. Across the cortex, there was a mean reduction of 0.56% (SD: 0.26 range: −1.10 to 0.22) in thickness per Tanner stage (see Figure 4b). Refer to Table S10 for model comparisons, and Table S11 for summary statistics of the linear model. An illustration of effect sizes by sex are presented in Figure S9.

**Figure 4.**
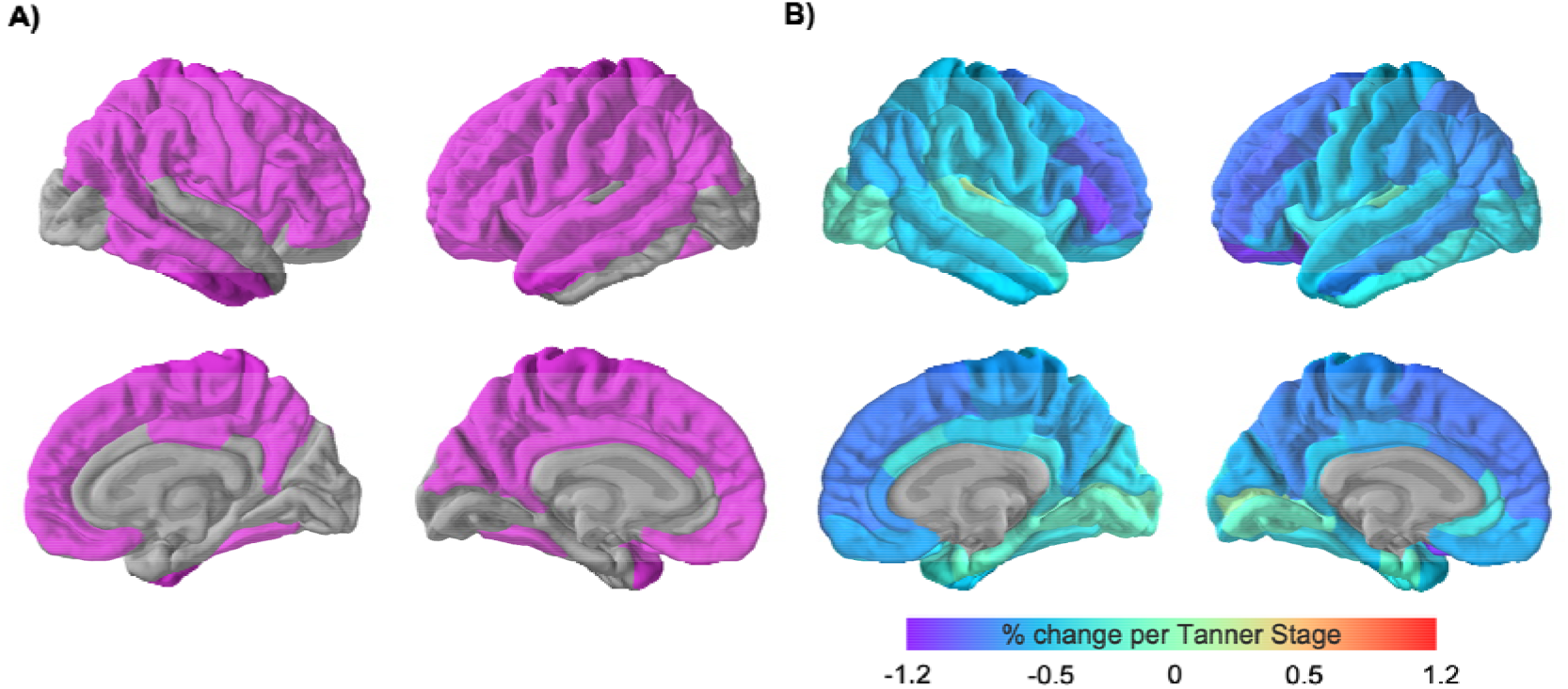
Puberty-related changes in cortical thickness, controlling for age. Regions exhibiting significant (linear) trajectories based on model comparisons (A). Effect sizes are illustrated as percent change per Tanner stage, relative to the average thickness at wave 1 (B). Refer to figure S7 for a comparison of effect sizes with and without controlling for age.

### Cortical development & pubertal tempo

Change in Tanner stage with age was modelled separately in males and females. Females exhibited linear increases in Tanner stage over time, while males exhibited a quadratic trajectory (see Figure 5 and Table S12). Models included a random intercept and (linear) age-slope, which were significantly correlated with one another in both females (0.58, p < 0.001) and males (0.64, p < 0.001). Random slopes from these models were used as an index of pubertal tempo in subsequent analyses.

**Figure 5.**
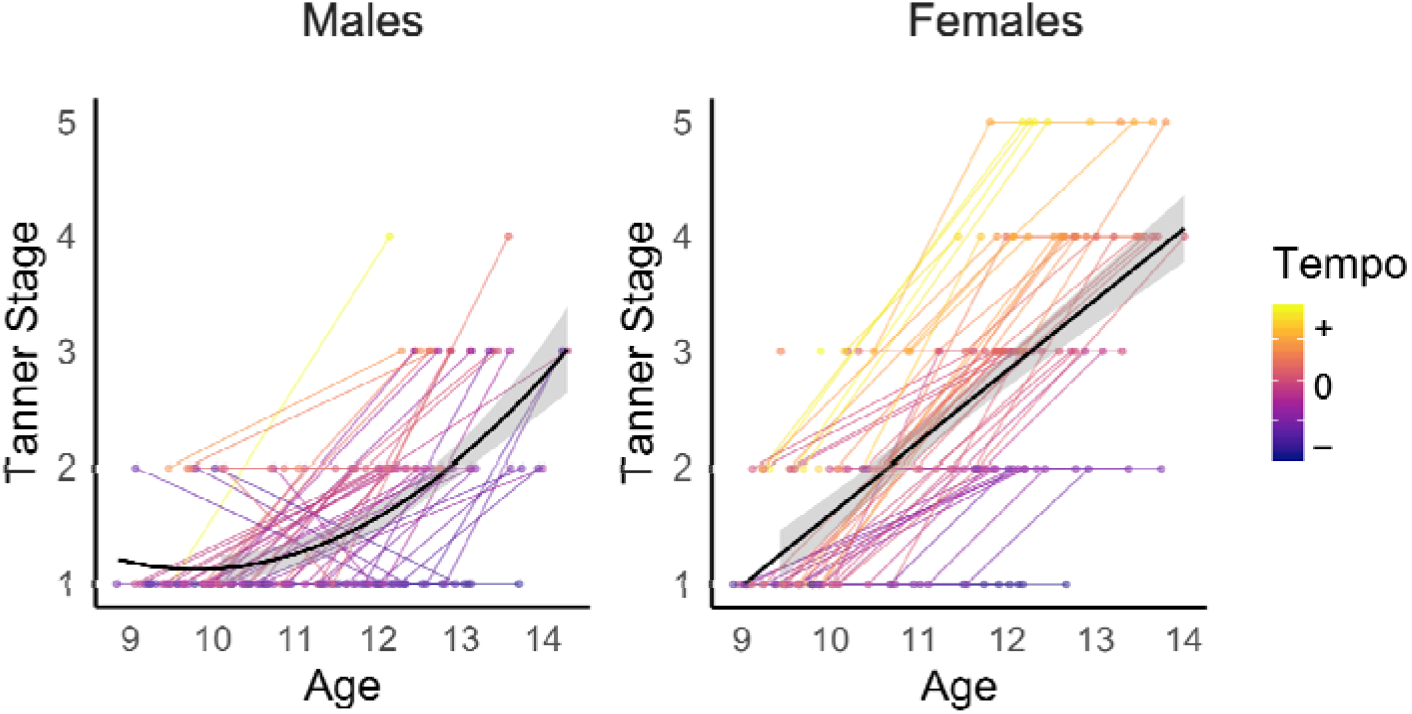
Changes in Tanner stage with age, modelled separately in males and females. Tempo: pubertal tempo, as indexed by random slopes from linear mixed models.

Interactions between pubertal tempo (i.e., random slopes) and age in relation to cortical thickness were examined within each sex. In males, this interaction improved model fit over and above age-related trajectories and a main effect of tempo in multiple lateral and medial frontal cortices, as well as bilateral precuneus, and left superior parietal and right middle temporal cortices (Figure 6A, with results presented in Tables S13 and S14). These regions also exhibited significant interaction terms for age and tempo (i.e., “s(age, by = tempo, bs = “cs”, k = 3)”) following FDR correction for multiple comparisons. As illustrated for the left caudal middle frontal cortex in Figure 6B, males with faster tempo exhibited greater reductions in thickness over time relative to those with slower tempo. Comparatively, there were no significant interactions between pubertal tempo and age in relation to cortical thickness in females (see Tables S15 and S16).

**Figure 6.**
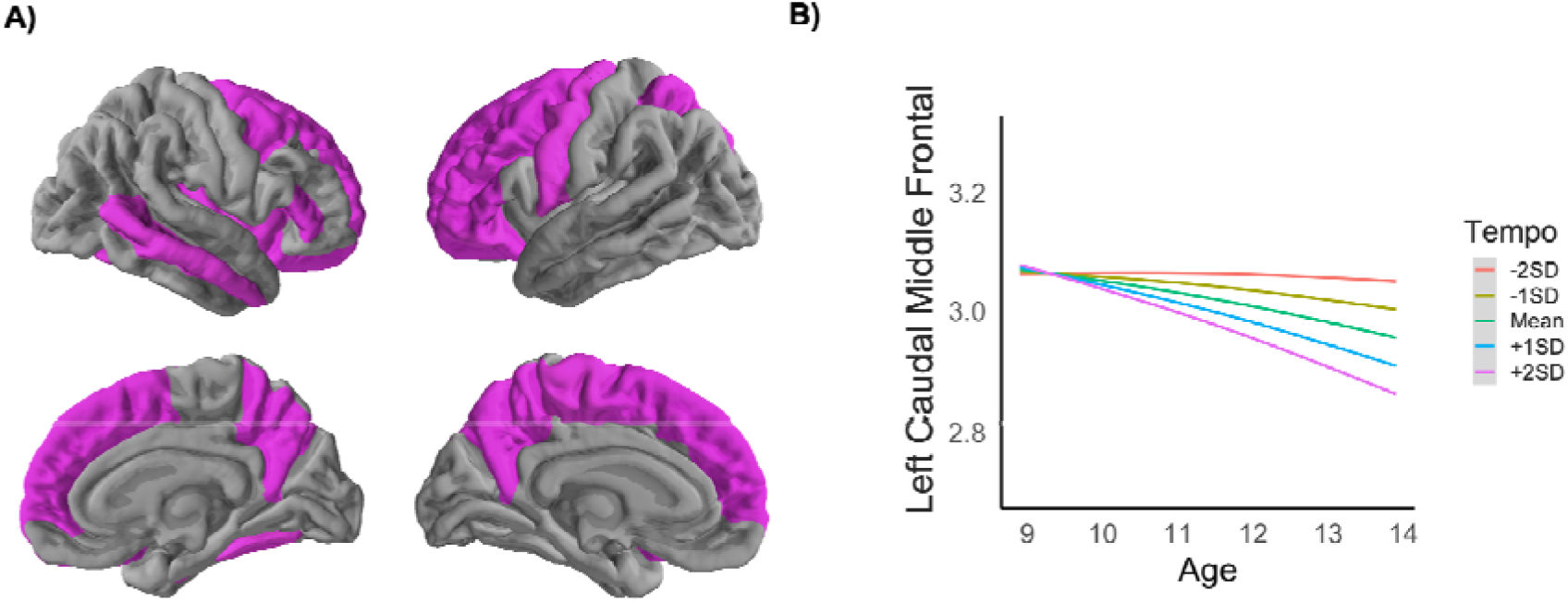
Variability in cortical development as a function of pubertal tempo in males (A), with effects illustrated for the left caudal middle frontal cortex (B).

### Sensitivity analyses

Sensitivity analyses were conducted *i)* by controlling for body-mass index and SES, *ii)* with the exclusion of participants on steroid medications over the year prior to the assessment, and *iii)* with the exclusion of participants who exhibited reductions in Tanner stage over time. Across these analyses, a few of the significant results became weaker and non-significant. However, the majority of results, and the overall pattern of results, remained consistent (see Tables S6, S10, S13 and S15).

## Discussion

This longitudinal investigation revealed extensive reductions in cortical thickness with pubertal maturation. These reductions were largely linear, and mostly consistent across males and females. These normative (group-level) changes explained additional variance beyond age-related cortical trajectories, suggesting that pubertal processes have a unique influence on structural brain development across adolescence. Moreover, in males, individual differences in the rate of pubertal maturation (or “pubertal tempo”) were associated with variation in cortical developmental trajectories in the frontal and precuneus cortices between late childhood and mid-adolescence.

We identified reductions in cortical thickness as individuals progressed through pubertal stages. These normative group-level trajectories survived correction for multiple comparisons across the entire cortex, apart from the bilateral entorhinal cortices. The pattern of reductions was found to be linear across the cortex, with regions exhibiting up to 2% change per pubertal stage. The strongest developmental effects were present within the left lateral prefrontal, superior parietal, and medial posterior (i.e., cuneus and precuneus) cortices. Most regions (particularly within the frontal and parietal cortices) also exhibited significant puberty-related changes when controlling for age, suggesting that pubertal stage may uniquely influence cortical development independent of age effects. Interestingly, controlling for age resulted in the strongest pubertal associations in the lateral prefrontal cortex (with regions exhibiting up to 1.2% change per pubertal stage) and considerably decreased the strength of pubertal associations in parts of the occipital and temporal cortices. The inclusion of age as a covariate in these models is akin to using residual pubertal stage (from Tanner stage ~ age) and may thus be interpreted as pubertal timing in some circumstances (i.e., individual variance beyond group-level associations between age and pubertal stage). However, such a residual pubertal variable was confounded by Tanner stage in our sample, as we did not have a representation of individuals with lower residual scores at higher Tanner stages given the age span (see Figure S10). In other words, higher residuals were correlated with higher Tanner stage. As such, our interpretation of these findings focuses on the unique role of Tanner stage beyond age, and not the potential role of pubertal timing.

Overall, these findings are largely consistent with prior cross-sectional studies that have identified negative associations between pubertal stage and thickness of the prefrontal and parietal cortices (Koolschijn et al., 2014; Peper et al., 2009). However, our results are more widespread across the cortical surface than hypothesized, which likely reflects the increased power to identify development processes with longitudinal data (relative to prior cross-sectional literature), as well as the ability to distinguish between age and pubertal process. Systematic reviews have also indicated distributed cortical associations with pubertal sex hormones (testosterone and estradiol; see Figure 1 in Vijayakumar et al., 2018), which may be contributing to the extent of current associations with pubertal stage – a global measure of underlying hormonal processes. Overall, these results are also suggestive of a role of puberty in the continued development of high-order cognitive and socioemotional processes that are supported by the frontoparietal cortices, highlighting the need for future research to consider pubertal maturity when investigating the neurodevelopmental correlates of adolescent behavior and functioning.

These patterns of puberty-related cortical development were consistent across males and females, with exploratory analyses failing to identify sex differences (with and without controlling for age-related changes, although effect sizes were larger in males, particularly when controlling for age - see figures S8 and S9). However, the underlying neuroendocrine mechanisms are likely to differ across sex. Neural changes are more likely to be driven by adrenarcheal processes in males given the earlier (and limited) window of pubertal maturation that was examined (as illustrated in Figure 2), suggesting the adrenal hormones (i.e., DHEA, DHEA-S) may play an important role in the identified patterns of cortical thinning. Comparatively, both adrenarcheal and gonadarcheal processes may be implicated in females as sampling incorporated the later stages of puberty, and as such, estradiol may also contribute to the identified patterns of cortical development.

The lack of sex differences in our exploratory analyses was unexpected as animal research suggests that sexual dimorphism in brain development may be attributed to differences in exposure to pubertal sex hormones, the density of hormone receptors, and cell-intrinsic mechanisms (for a review, see Denley et al., 2018; Juraska & Willing, 2017). Most animal studies have focused on limbic regions (e.g., amygdala, hippocampus, and hypothalamus; Ahmed et al., 2008; De Lorme et al., 2012), and corresponding sex differences have been identified in longitudinal human research on subcortical development, particularly in relation to pubertal hormone concentrations (Goddings et al., 2014; Herting et al., 2014).

Nevertheless, animal research has also implicated pubertal processes in sexual dimorphism in prefrontal and occipital cortical development (Antonio Muñoz-Cueto et al., 1990; Koss et al., 2015; Markham et al., 2007) that may lead us to expect sex differences in puberty-related development of these cortical regions. However, the lack of findings could be attributed to females being overall more pubertally advanced than males in the current dataset. An alternate consideration is that Tanner stage may lack the specificity that is required to identify sex differences. Thus, to gain a better understanding of potential sex differences in puberty-related cortical development, further research is needed that *i)* covers equivalent periods of pubertal development in males and females, and *ii)* considers hormonal processes.

Beyond normative (i.e., group-level) cortical development, we identified variability in cortical developmental trajectories as a function of pubertal tempo in males. Inconsistent with our hypothesis, males with faster pubertal tempo (i.e., those who were quicker to progress through the Tanner stages relative to their peers) exhibited greater reductions in cortical thickness (or greater acceleration of cortical thinning) relative to those who exhibited slower pubertal tempo. This pattern of development was identified for a number of bilateral prefrontal cortices (specifically the caudal middle frontal, superior frontal, lateral orbitofrontal and parstriangularis) and the precuneus, as well as the left rostral middle frontal, pars orbitalis, and superior parietal, and right middle temporal cortices. As there was little variance in pubertal stage at baseline for males, these associations were specific to temporal change in pubertal maturity. Although Herting and colleagues (2015) found that greater increases in pubertal stage (over a 2-year period) was related to less thinning of the right superior frontal and superior temporal cortices in 10 to 16 year olds (across sex), they also found that greater increases in estradiol concentrations were related to greater thinning in the left middle temporal cortex (in females). Moreover, Wierenga and colleagues (2018) showed that adolescents with earlier pubertal timing (indicative of more advanced pubertal stage at a given point in time) had larger subcortical structures than their peers, suggestive of accelerated patterns of normative development in these regions. Thus, we speculate that our findings may also be reflective of earlier or accelerated cortical maturation in male adolescents with faster pubertal tempo.

Taken together, our analyses show that most of the cortex exhibits thinning as individuals progress through the stages of puberty, but specific regions that cluster primarily in the frontal and parietal cortices also exhibit accelerated development in males who are quicker to progress through puberty relative to peers. The implicated frontoparietal regions also subserve a number of goal-directed cognitive and affective functions, including the regulation of attention, executive functions, inhibition of actions, and self-referential processes (Cole et al., 2014; Scolari et al., 2015). Thus, pubertal tempo-related cortical development may specifically indicate alterations in the maturation of these cognitive and affective processes in males. Indeed, it is unlikely that normative changes in brain structure during puberty will account for variation in cognition, affect or behavior; rather individual differences in puberty and related cortical development are more likely to do so. However, it remains unclear whether earlier or accelerated cortical development is adaptive. Earlier brain development could suggest that individuals have greater cognitive capacities as they are quicker to acquire adult levels of maturity (Shaw et al., 2006). Conversely, earlier development could result in a mis-match between biological processes and the environment, which may increase risk for psychopathology (Ullsperger & Nikolas, 2017). This would be consistent with prior research that has found faster pubertal tempo to be related to greater internalizing and externalizing symptoms in males (Beltz et al., 2014; Marceau et al., 2011; Mendle et al., 2010). Thus, it is important that future research relate variability in puberty-related cortical development with cognition and affect in order to improve our understanding of how these biological processes influence behaviour and functioning during the transition from childhood to adolescence.

We speculate that sex differences in pubertal development in the dataset may also contribute to the lack of associations between pubertal tempo and cortical development in females. While linear rates of tempo may appropriately capture the earlier (and limited) window of pubertal maturation in males, nonlinear trajectories may be required to study the more extended period in females. Indeed, prior work has identified such nonlinear patterns of (individual-level) pubertal changes, and found these trajectories to relate to socioemotional functioning in females (Marceau et al., 2011). Although variability in nonlinear pubertal development (i.e., random slopes) were not modelled in females here, future studies with more than three time points are needed to estimate such complex individual trajectories, and gain a better understanding of the full pubertal period.

### Strengths & limitations

The strengths of this study include the use of a large sample of late childhood to adolescence, as well as a flexible modelling strategy of repeated neuroimaging assessments over multiple time points across pubertal stages. We specifically increased sample size (and thus power) by combining two longitudinal cohorts. We modelled cortical changes using generalized additive mixture models that advance on traditional linear models as they i) do not impose a specific developmental trajectory, and ii) fit varied trajectories to different groups, thus allowing us to examine for sex differences in both the shape and slope of trajectories.

Despite these strengths, our findings need to be considered within the context of some limitations. Variance in pubertal maturation at the later stages was limited, particularly in males. Thus, examination of sex differences was exploratory, and the lack of significant sex differences could result from the lack of later stages in males. Moreover, future research may benefit from recruiting males that are 1 to 2 years older than females, as undertaken in some previous work at the University of Pittsburgh (Herting et al., 2015). Extended assessments into late adolescence (i.e., 17 to 18 years) is also required to capture the full spectrum of pubertal maturation across individuals, and to better understand how these patterns relate to later socioemotional outcomes. While the NICAP cohort was assessed on an upgraded scanner at the last wave of assessments, this upgrade was not confounded with pubertal development (see Figure S3). We were therefore able to account for potential scanner differences in our statistical models.

Our analyses focused on parent report of pubertal stage, which was primarily completed by mothers (94% of respondents). Parent-report has been shown to correspond well to clinician ratings in females (Brooks-Gunn et al., 1987; Huang et al., 2012), but less so in males (Huang et al., 2012). Thus it is possible that limitations with the parent-report Sexual Maturation Scale may be contributing to sex-specific findings. However, it is important to note that selfreport also has limitations; adolescents at lower and upper pubertal stages tend to report towards the mid-stages (Huang et al., 2012; Shirtcliff et al., 2009), and there are sex differences in correspondence between self-report and pubertal hormone concentrations (Shirtcliff et al., 2009). Moreover, parent- and self-report relate similarly to hormone concentrations (Huang et al., 2012). Given that both questionnaire assessments face limitations (but are also more feasible that clinician ratings in such cohort studies), future work may benefit from collecting both measures and checking correspondence in cortical trajectories when using parent- and self-report. Pubertal stage is also a gross measure of a number of underlying biological processes, and future studies may provide further insight by considering measures of specific pubertal processes, such as hormone concentrations and expression of steroid hormone receptors in the brain. We were also unable to examine the effects of pubertal timing. As discussed above, the residual timing index was confounded by Tanner stage in our sample, as we did not have a representation of adolescents with lower residual scores (i.e., “later timing”) at higher Tanner stages given the age span. Moreover, we were unable to measure timing according to age of pubertal onset with precision as the time interval between assessments ranged up to 3.3 years, but this represents an important measure for future research to identify neurodevelopmental trajectories that may represent risk for future socioemotional problems. Finally, we were unable to control for race and ethnicity as this data was not collected for one of the examined cohorts. Given the well-established race/ethnic disparities in age of pubertal onset (Biro et al., 2010; Chumlea et al., 2003; Hoyt et al., 2018; Wu et al., 2002), this represents an important consideration for future research.

### Conclusions

The current longitudinal investigation highlights widespread puberty-related cortical development, including thinning of a number of regions that support higher order cognitive processes, over and above well-documented age-related changes in the cortex. Importantly, these findings suggest that the unique influence of puberty on brain development may be more extensive than previously identified. Moreover, variation in cortical trajectories with pubertal tempo (particularly within the prefrontal cortex in males) may relate to behavioral and emotional functioning during adolescence. Further work is needed to investigate the implications of the current findings for understanding the biological risk and resilience mechanisms for socioemotional problems in adolescents.

## Acknowledgements

We would like to thank all of the families who have participated in both NICAP and iCATS. NICAP was funded by the National Medical Health and Research Council of Australia (NHMRC; project grant #1065895). iCATS was funded by a Discovery Project grant from the Australian Research Council (ARC; DP120101402). SW was supported by a NHMRC Career Development Fellowship (#1125504). Cohorts were supported by the Murdoch Children’s Research Institute, The Royal Children’s Hospital, The Royal Children’s Hospital Foundation, Department of Paediatrics at The University of Melbourne and the Victorian Government’s Operational Infrastructure Support Program. We also thank The Royal Children’s Hospital’s Medical Imaging staff for their assistance and expertise in the collection of the MRI data included in this study.

