## Supplementary Materials for "A longitudinal analysis of puberty-related cortical development"

**Supplementary Material**

**NICAP:** Following exclusion for poor quality structural MRI (see below for further detail), the final dataset consisted of 207 (114 male) observations across the three waves from 90 adolescents, specifically comprising 14 (8 male) adolescents with one observation, 35 (20 male) with any (and only) two observations, and 41 (22 male) with all three observations. The age ranges and sex distribution at each time point of the control participants are reported in Table S1. Attrition analyses, presented in Table S2, did not reveal any significant differences in age, pubertal stage, sex, or SES in adolescents who completed the final wave of assessments to those who did not.

Table S1. Sex distribution and age at each wave for NICAP

|  | Wave 1 | Wave 2 | Wave 3 |
| --- | --- | --- | --- |
| *Male* |  |  |  |
| N | 47 | 39 | 28 |
| Age | 10.46, 0.50, 9.58-11.86 | 11.77, 0.56, 10.88-13.45 | 13.25, 0.58, 12.2-14.29 |
| *Female* |  |  |  |
| N | 35 | 36 | 22 |
| Age | 10.39, 0.34, 9.86-11.02 | 11.74, 0.38, 11.02-12.9 | 13.25, 0.44, 12.43-14.0 |

Note: Age reports mean, SD, range (years)

Table S2. Attrition analyses for NICAP – differences at baseline (wave1).

|  | Missing | Not missing |  |
| --- | --- | --- | --- |
| Age | 10.52 | 10.35 | t(75) = -1.863, p = 0.066 |
| Sex (M:F) | (22:18) | (28:22) | X^2^(1) = 0, p = 1 |
| Tanner stage | 1.34 | 1.30 | t(72) = -0.408, p = 0.685 |
| SES | 1014 | 1021 | t(76) = 0.669, p = 0.506 |

Note: Groups (missing vs not missing) are categorized based on inclusion in study at wave 3.

**iCATS:** Following exclusion for poor quality structural MRI (see below for further detail), the final dataset consisted of 159 (72 male) observations across the two waves from 102 adolescents, specifically comprising 45 (20 male) adolescents with one observation and 57 (26 male) with two observations. The age ranges and sex distribution at each time point of the control participants are reported in Table S3. Attrition analyses, presented in Table S4, did not reveal any significant differences in age, pubertal stage, sex, or SES in adolescents who completed the final wave of assessments compared to those who did not.

Table S3. Sex distribution and age at each wave for iCATS

|  | Wave 1 | Wave 2 |
| --- | --- | --- |
| *Male* |  |  |
| N | 35 | 37 |
| Age | 9.62, 0.35, 8.85-10.25 | 12.42, 0.37, 11.81-13.26 |
| *Female* |  |  |
| N | 44 | 43 |
| Age | 9.50, 0.32, 8.84-10.19 | 12.22, 0.41, 11.51-13.30 |

Note: Age reports mean, SD, range (years)

Table S4. Attrition analyses for iCATS – differences at baseline (wave1).

|  | Missing | Not missing |  |
| --- | --- | --- | --- |
| Age | 9.62 | 9.54 | t(32) = -0.950, p = 0.349 |
| Sex (M:F) | (9:13) | (37:43) | X^2^(1) = 0.042, p = 0.838 |
| Tanner stage | 1.30 | 1.28 | t(25) = -0.147, p = 0.885 |
| SES | 1063 | 1054 | t(34) = -0.584, p = 0.563 |

Note: Groups (missing vs not missing) are categorized based on inclusion in study at wave 2.

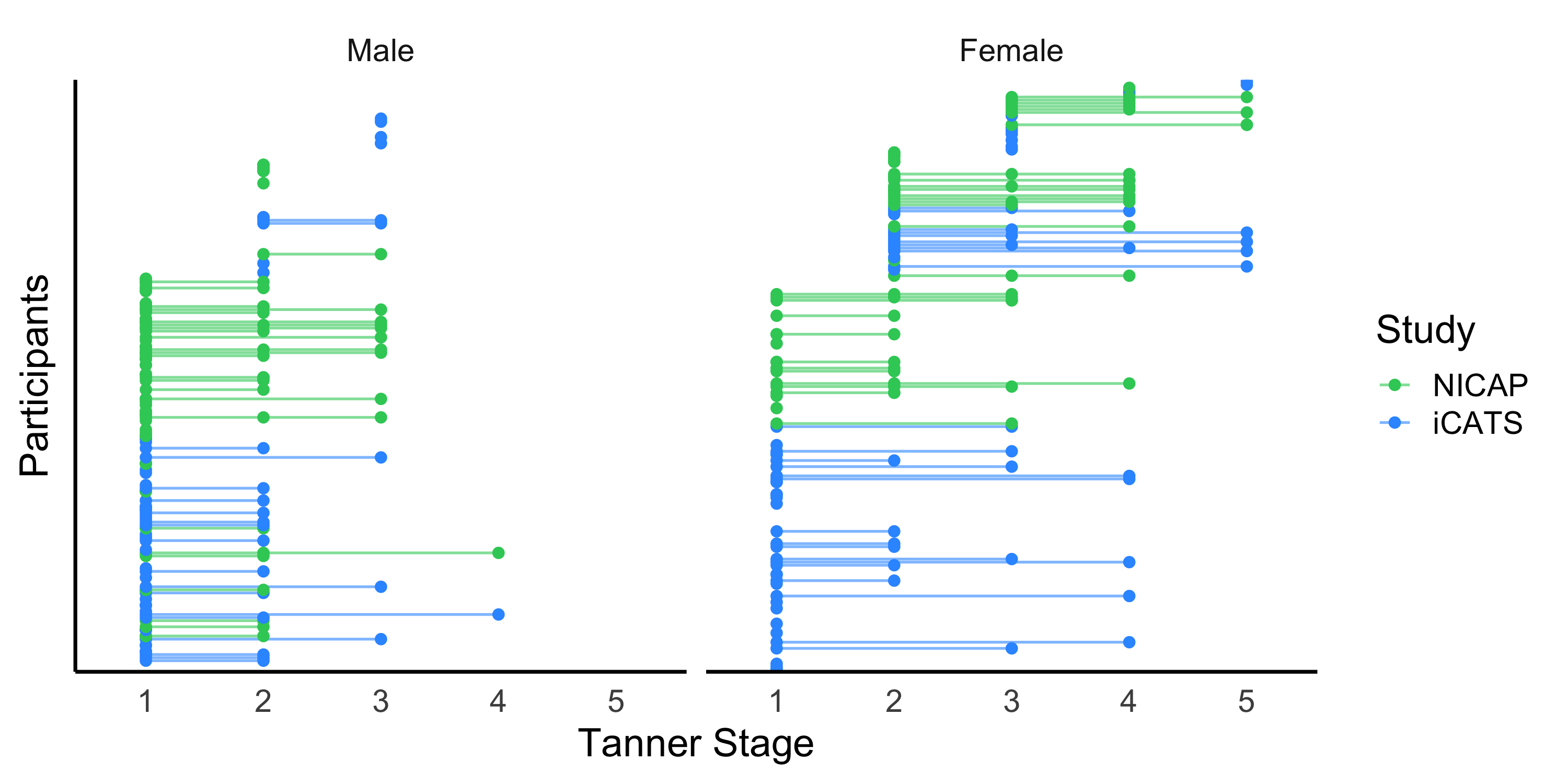
*Figure S1.* Distribution of pubertal (Tanner) stage for each cohort

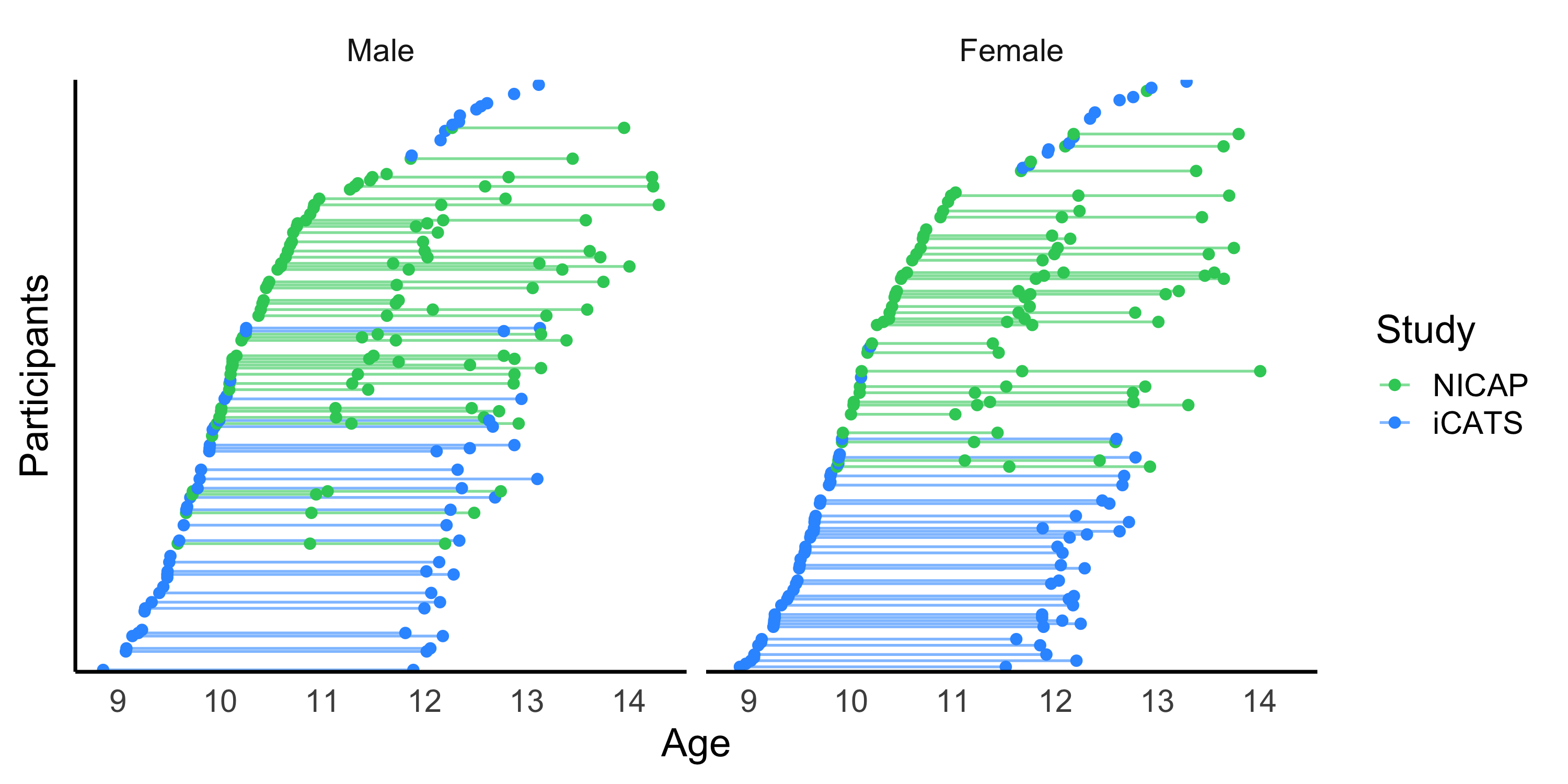

*Figure S2.* Distribution of age for each cohort

**NICAP Quality Control**

***Manual QC.*** The quality of the raw and processed T1-weighted images was visually inspected. Two independent raters were trained on quality assessment of the raw images, and another two raters were trained on quality assessment of the FreeSurfer reconstruction (i.e., processed images). All raters were trained using a quality control manual created by a senior researcher, and reliability of each rater was assessed after the first 50 images (approximately 10% of the dataset).

The raw images were rated on a 4-point scale for two scales: “ringing” and “blurriness”. Scores of “1” reflected no ringing and sharply defined images, “2” reflected slight ringing that was restricted to a small (cortical) area or slight blurriness, “3” reflected more ringing that extended deeper into white matter and covered more brain regions or considerable blurriness, and “4” reflected extensive ringing and blurring throughout the brain. Both raters checked the full dataset, compared scoring discrepancies for each scale, and decided on a consensus rating for these images. Images with a final score of “3” or greater on either subscale were excluded (N=9, male=7).

The processed images were rated on a 3-point scale, focusing on the accuracy of the white and pial surfaces. Scores of “1” reflected near perfect reconstruction, “2” reflected minor reconstruction issues that were limited to small areas of the brain, and “3” reflected poor reconstruction (e.g., consistent under-estimation of white matter, or extensive areas where CSF was included as grey matter). Again, both raters checked the full dataset, compared scoring discrepancies, and decided on a consensus rating for these images. Images with a final score of “3” were excluded. This process identified a further 3 images (male=2) that were excluded following QC of raw images.

***Automated QC.*** We also processed all raw T1 images through MRIQC (v0.14.2), to supplement the manual QC. MRIQC provides automated prediction of the quality of MRI scans, including a binary classifier that labels images as include vs. exclude. The classifier, which has been trained on publicly available multi-site datasets (17 sites), flagged 1 image for exclusion (which had not been excluded in the manual QC assessments). Further visual inspection of this image by a senior researcher deemed that it should be excluded.

***QC exclusions.*** The final excluded NICAP sample from manual and automated QC thus consisted of a total of 13 scans from 12 participants (males = 9).

**iCATS Quality Control**

***Manual QC.*** The quality of the raw and processed T1-weighted images was visually inspected by the same raters, and using the same rating scale, as the NICAP sample.

Two independent raters checked the quality of the raw images, and both raters checked the full dataset. A senior researcher subsequently checked images that were scored a “3” or “4” on either scale by either rater, and made a final decision regarding quality. Images with a final score of “3” or “4” on either scale were excluded (N=24, males=12), leaving a sample of 159 images that were processed through FreeSurfer.

The 159 processed images were rated on a 3-point scale, focusing on the accuracy of the white and pial surfaces. Scores of “1” reflected near perfect reconstruction, “2” reflected minor reconstruction issues that were limited to small areas of the brain, and “3” reflected poor reconstruction (e.g., consistent under-estimation of white matter, or extensive areas where CSF was included as grey matter). Two raters were trained on quality assessment of the FreeSurfer reconstruction (i.e., processed images). The first rater checked the entire dataset, and the second rater subsequently checked all images that were scored a “2” or “3” (N = 66) and a subset of images that were scored a “1” (N = 32, 33%). Finally, a senior researcher checked images that were scored a “3” by either rater, and made a final decision regarding quality. Images with a final score of “3” were excluded. This did not identify any further images that were excluded following QC of raw images, thus leaving a sample of 159 images.

***Automated QC.*** We also processed all raw T1 images through MRIQC (v0.14.2), to supplement the manual QC. MRIQC provides automated prediction of the quality of MRI scans, including a binary classifier that labels images as include vs. exclude. The classifier. which has been trained on publicly available multi-site datasets (17 sites), flagged 9 images for exclusion. 6 of these images overlapped with exclusions determined by manual QC assessments. Further visual QC by a senior researcher deemed that the remaining 3 images were of sufficient quality to be included in analyses.

***QC exclusions.*** The final excluded iCATS sample from manual and automated QC thus consisted of a total of 24 scans from 22 participants (males = 12).

**Scanner upgrade**

Wave 3 of the NICAP MRI assessment was undertaken on a different (upgraded) scanner to the previous waves of NICAP and both waves of iCATS. The distribution of scanner upgrade as a function of pubertal stage and age is illustrated in Figures S3 and S4, respectively.

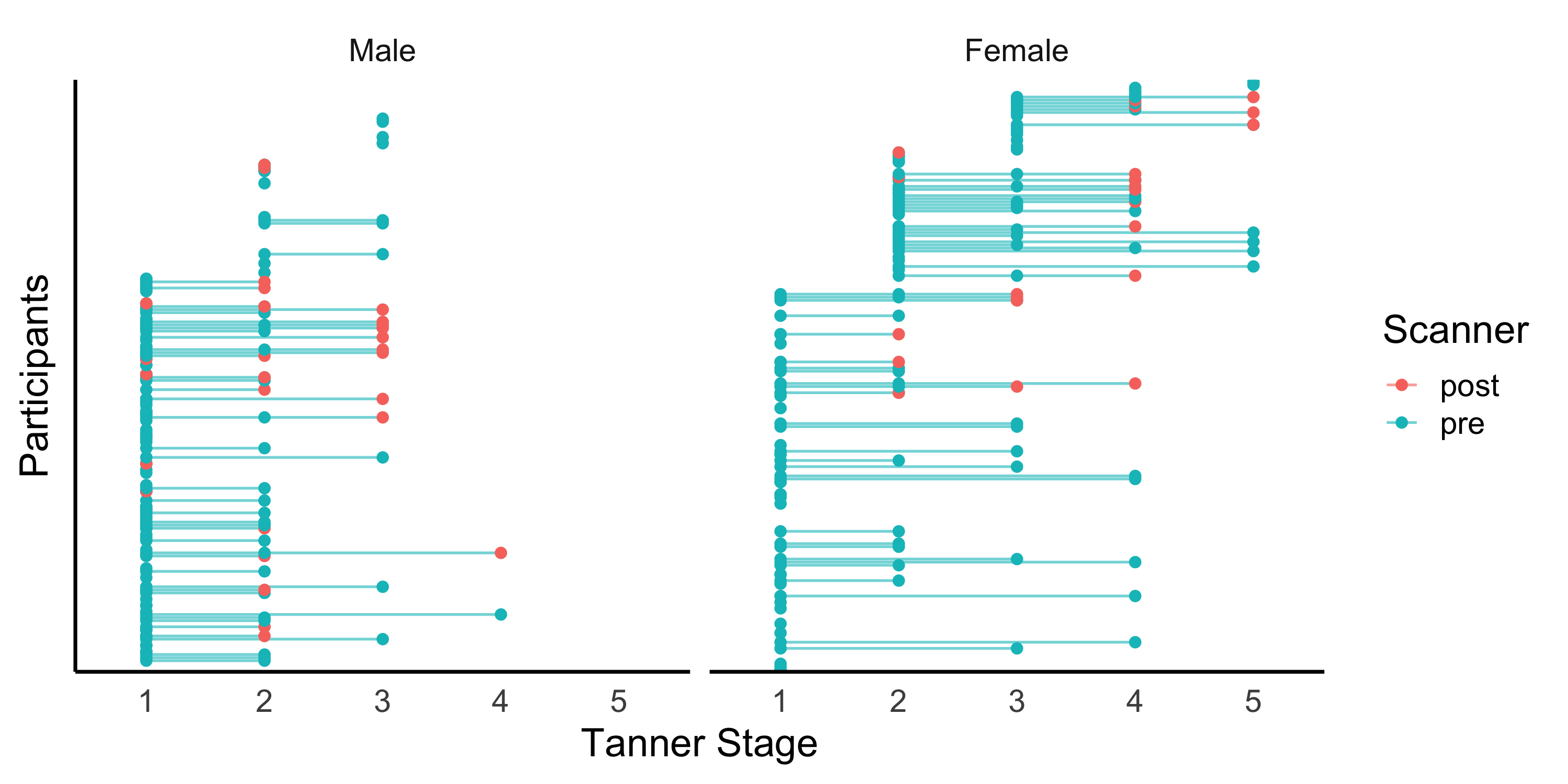
*Figure S3.* Distribution of scanner differences by pubertal (Tanner) stage

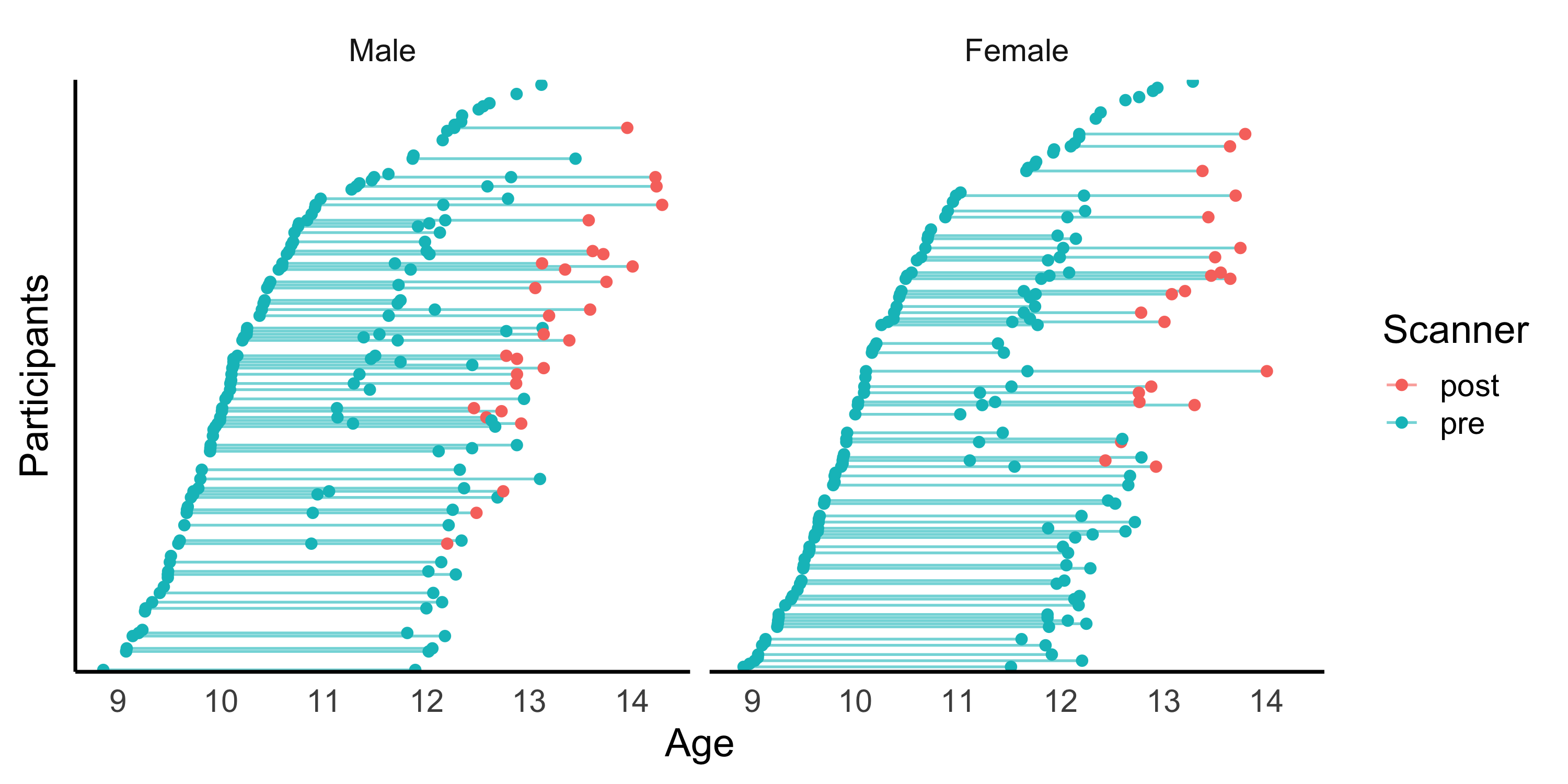
*Figure S4.* Distribution of scanner differences by age

In order to examine the potential effects of the scanner upgrade of cortical thickness estimates, we compared brain structure a subsample of age-matched (and sex-matched) NICAP participants who completed scans prior to the scanner upgrade (“pre”; N = 11, 72% males) and following the scanner upgrade (“post”; N = 11, 72% males). All participants were between 12.13 and 13.45 years of age. We compared cortical thickness of the Desikan–Killiany–Tourville parcellation between the “pre” and “post” groups, with the hypothesis that potential scanner differences would present as significant group differences after controlling for age and sex. We examined this hypothesis with a series of linear regressions predicting thickness of each region from group. The right insula was the only region that exhibited significant group differences in cortical thickness following FDR correction, however a few other regions exhibited differences that did not survive correction for multiple comparisons (left pars opercularis, left pars orbitalis, left insula, left precentral, left superior temporal, right superior temporal, right inferior parietal, right lateral orbitofrontal). Across these regions, there was significantly greater cortical thickness in the group of adolescents that were scanned post scanner upgrade (see Table S5).

In order to account for the scanner upgrade, a binary scanner variable (i.e., “pre-upgrade” or “post-upgrade”) was incorporated into statistical models. Given the spread of post-upgrade scans across pubertal stage (see Figure S3), we did not expect the puberty-related results to be confounded by the scanner upgrade. To ensure that age-related models (presented in the Supplement) were not confounded by scanner differences, we additionally ran analyses excluding participants over 13 years of age. This subsample (N = 329, with 18 participants scanned post-upgrade) exhibited a similar rate of cortical change to the full sample (see Figure S6).

Table S5. Cortical thickness differences between “pre” and “post” scanner upgrade groups.

| Region | B | SE | p | fdr-corrected |
| --- | --- | --- | --- | --- |
| right insula | 0.268 | 0.045 | 0 | 0 |
| left pars opercularis | 0.16 | 0.057 | 0.011 | 0.124 |
| left pars orbitalis | 0.258 | 0.078 | 0.004 | 0.124 |
| left insula | 0.178 | 0.057 | 0.006 | 0.124 |
| right inferior parietal | 0.148 | 0.052 | 0.011 | 0.124 |
| right lateral orbitofrontal | 0.123 | 0.044 | 0.012 | 0.124 |
| right superior temporal | 0.113 | 0.042 | 0.014 | 0.124 |
| left superior temporal | 0.123 | 0.047 | 0.017 | 0.132 |
| left precentral | 0.12 | 0.047 | 0.021 | 0.145 |
| right middle temporal | 0.117 | 0.057 | 0.053 | 0.244 |
| right pars opercularis | 0.145 | 0.067 | 0.045 | 0.244 |
| right pars triangularis | 0.143 | 0.068 | 0.05 | 0.244 |
| right postcentral | 0.088 | 0.042 | 0.049 | 0.244 |
| right precentral | 0.106 | 0.052 | 0.055 | 0.244 |
| left posterior cingulate | 0.1 | 0.053 | 0.073 | 0.251 |
| right inferior temporal | 0.081 | 0.043 | 0.072 | 0.251 |
| right lateral occipital | 0.094 | 0.049 | 0.069 | 0.251 |
| right supramarginal | 0.101 | 0.052 | 0.066 | 0.251 |
| left superior frontal | 0.087 | 0.047 | 0.08 | 0.261 |
| left pars triangularis | 0.139 | 0.076 | 0.086 | 0.267 |
| right lingual | 0.117 | 0.066 | 0.094 | 0.268 |
| right parsorbitalis | 0.133 | 0.076 | 0.095 | 0.268 |
| left lingual | 0.089 | 0.053 | 0.106 | 0.286 |
| left middle temporal | 0.092 | 0.056 | 0.115 | 0.297 |
| left inferior parietal | 0.079 | 0.049 | 0.121 | 0.3 |
| left caudal anterior cingulate | 0.121 | 0.079 | 0.14 | 0.308 |
| left caudal middle frontal | 0.065 | 0.043 | 0.149 | 0.308 |
| left postcentral | 0.085 | 0.054 | 0.132 | 0.308 |
| right entorhinal | -0.155 | 0.099 | 0.136 | 0.308 |
| right isthmus cingulate | 0.093 | 0.062 | 0.154 | 0.308 |
| right paracentral | 0.116 | 0.077 | 0.149 | 0.308 |
| left entorhinal | -0.165 | 0.115 | 0.167 | 0.324 |
| right medial orbitofrontal | 0.127 | 0.094 | 0.191 | 0.356 |
| right rostral anterior cingulate | 0.109 | 0.081 | 0.195 | 0.356 |
| left precuneus | 0.062 | 0.048 | 0.21 | 0.361 |
| left supramarginal | 0.069 | 0.054 | 0.22 | 0.361 |
| right rostral middle frontal | 0.087 | 0.068 | 0.218 | 0.361 |
| right superior frontal | 0.065 | 0.051 | 0.221 | 0.361 |
| left transverse temporal | 0.097 | 0.082 | 0.248 | 0.394 |
| left lateral occipital | 0.053 | 0.046 | 0.257 | 0.398 |
| right posterior cingulate | 0.055 | 0.049 | 0.274 | 0.414 |
| right parahippocampal | -0.087 | 0.085 | 0.322 | 0.475 |
| left isthmus cingulate | 0.069 | 0.072 | 0.347 | 0.5 |
| left paracentral | 0.054 | 0.065 | 0.415 | 0.541 |
| left rostral anterior cingulate | 0.075 | 0.088 | 0.402 | 0.541 |
| left rostral middle frontal | 0.053 | 0.064 | 0.419 | 0.541 |
| right cuneus | 0.053 | 0.063 | 0.414 | 0.541 |
| right precuneus | 0.04 | 0.049 | 0.417 | 0.541 |
| right transverse temporal | 0.091 | 0.118 | 0.449 | 0.568 |
| left medial orbitofrontal | 0.04 | 0.054 | 0.475 | 0.587 |
| right caudal middle frontal | 0.045 | 0.062 | 0.483 | 0.587 |
| left inferior temporal | 0.028 | 0.045 | 0.539 | 0.608 |
| left lateral orbitofrontal | 0.039 | 0.062 | 0.538 | 0.608 |
| left pericalcarine | 0.045 | 0.072 | 0.537 | 0.608 |
| right pericalcarine | 0.054 | 0.085 | 0.532 | 0.608 |
| left fusiform | 0.018 | 0.038 | 0.643 | 0.712 |
| left cuneus | 0.019 | 0.057 | 0.745 | 0.802 |
| right fusiform | 0.015 | 0.046 | 0.75 | 0.802 |
| right caudal anterior cingulate | -0.015 | 0.059 | 0.804 | 0.845 |
| left parahippocampal | 0.015 | 0.112 | 0.894 | 0.924 |
| left superior parietal | 0.002 | 0.057 | 0.969 | 0.984 |
| right superior parietal | 0.001 | 0.051 | 0.984 | 0.984 |

**Imputation**

Tanner stage was missing for 26 data points in NICAP (12%, males = 15) and 7 data points in iCATS (4%, males = 5). Multiple imputation was undertaken using “mice” (Buuren & Groothuis-Oudshoorn, 2011) in R (R Core Team, 2013) to deal with missing data. This was conducted separately for NICAP and iCATS, as different predictor variables were available to perform imputation in each cohort.

The imputation model for NICAP included sex, age and the parent-report Pubertal Development Scale (Petersen et al., 1988). The PDS consists of five questions that assess height growth, body hair and skin changes, as well as breast development and menarche in females, and facial hair and voice changes in males. Items are scored on a 4-point scale ranging from “no physical changes” to “development seems complete”, with the only exception being the yes/no item on menarche. We utilized the Shirtcliff, Dahl & Pollak (2009) method to score the PDS into a 5-point scale, in order to approximate Tanner staging.

The imputation model for iCATS included sex, age and pubertal hormone levels (specifically testosterone, dehydroepiandrosterone (DHEA) and DHEA-sulphate). Hormones were assayed in waking saliva samples. At wave 1, two saliva samples were collected: on the day prior to, and on the day of MRI scanning. At wave 2, two to four samples were collected with samples provided across consecutive weeks (including the day of scanning). For each sample, children collected 2.5 ml of saliva in a test tube, via passive drool. Participants recorded the time they started collection of saliva and the duration of collection using a stopwatch. Any samples from participants who indicated that samples were provided more than 45 min after waking were excluded. Samples were frozen at −30 °C, and prior to analysis, defrosted and centrifuged, with the supernatant assayed in duplicate for levels of DHEA, DHEA-S, and testosterone using Salimetrics ELISA kits. See Whittle et al. (2020) for inter- and intra-assay coefficients of variation. Analyses were conducted on averaged hormone concentrations, in order to minimize proximal confounds that can influence salivary biomarkers.

**Pubertal development**

Linear mixed models were used to examine cohort differences in pubertal development, within each sex. First, best fitting developmental models were identified by comparing linear and quadratic age effects to a null model (without age). All models controlled for cohort, and likelihood ratio tests were used to assess model fit. Next, the best-fitting development model was compared to a model including an interaction between age and cohort. Again, likelihood ratio tests were used to assess model fit. The results revealed linear changes in Tanner stage in females and quadratic changes in males. These trajectories did not differ across the two cohorts in either sex (i.e., model fit did not improve with the inclusion of the age-by-cohort interaction terms; see Table S11).

Next, linear mixed models examined whether there were sex differences in pubertal development. First, best fitting developmental models were identified by comparing linear and quadratic age effects to a null model (without age) across both sexes. All models controlled for cohort and sex, and likelihood ratio tests were used to assess model fit. Next, the best-fitting development model was compared to a model including an interaction between age and sex. Again, likelihood ratio tests were used to assess model fit. Results revealed significant nonlinear sex differences in pubertal development (see Table S11).

Table S11. Model fit statistics for Tanner stage models.

|  | Model | DF | AIC | logLik | Test | Chisq | ChiDF | Pval |
| --- | --- | --- | --- | --- | --- | --- | --- | --- |
| *Females* |  |  |  |  |  |  |  |  |
| Null | A | 4 | 560.32 | -276.16 |  |  |  |  |
| **Linear** | **B** | **5** | **421.14** | **-205.57** | **1 vs 2** | **141.18** | **1** | **< 0.001** |
| Quadratic | C | 6 | 420.60 | -204.30 | 3 vs 2 | 2.54 | 1 | 0.11 |
| Linear * Cohort | D | 6 | 423.11 | -205.56 | 4 vs 2 | 0.03 | 1 | 0.87 |
| *Males* |  |  |  |  |  |  |  |  |
| Null | A | 4 | 415.51 | -203.75 |  |  |  |  |
| Linear | B | 5 | 341.30 | -165.65 | 1 vs 2 | 76.21 | 1 | < 0.001 |
| **Quadratic** | **C** | **6** | **330.29** | **-159.15** | **3 vs 2** | **13.01** | **1** | **< 0.001** |
| Quadratic * Cohort | D | 8 | 332.81 | -158.41 | 4 vs 3 | 1.48 | 2 | 0.48 |
| *Sex differences* |  |  |  |  |  |  |  |  |
| Null | A | 5 | 1034.33 | -512.17 |  |  |  |  |
| Linear | B | 6 | 830.82 | -409.41 | 1 vs 2 | 205.51 | 1 | < 0.001 |
| Quadratic | C | 7 | 820.26 | -403.13 | 3 vs 2 | 12.56 | 1 | < 0.001 |
| **Quadratic * Sex** | **D** | **9** | **783.79** | **-382.89** | **4 vs 3** | **40.74** | **2** | **< 0.001** |

In order to calculate pubertal tempo, we incorporated random age slopes to the best fitting model within each sex (i.e., linear in females and quadratic in males). Table S12 presents the results of these final models. The random intercepts and slopes were significantly positively correlated with one another in both females (R = 0.61, p < 0.001) and males (R = 0.65, p < 0.001). However, there was almost no variance in random intercept for males (SD = 2.04 e-26). Figure S5 presents the correlation between in random intercept and slope in females.

Table S12. Linear mixed models predicting Tanner stage from age, separately in each sex.

|  |  | Random | | Fixed |  |  |
| --- | --- | --- | --- | --- | --- | --- |
|  |  | Std Dev | Std Err | Estimate | Std Err | z |
| Female | Random intercept | 0.414 | 0.784 |  |  |  |
|  | Random slope | 0.276 | 0.031 |  |  |  |
|  | Residual | 0.394 | 0.041 |  |  |  |
|  | Fixed intercept |  |  | 1.022 | 0.073 | 14.09* |
|  | Age |  |  | 0.586 | 0.390 | 14.99* |
| Male | Random intercept | 0.000 | 0.000 |  |  |  |
|  | Random slope | 0.169 | 0.020 |  |  |  |
|  | Residual | 0.405 | 0.030 |  |  |  |
|  | Fixed intercept |  |  | 1.355 | 0.100 | 13.62* |
|  | Age |  |  | -0.372 | 0.112 | -3.34* |
|  | Age^2^ |  |  | 0.143 | 0.025 | 13.62* |

* p < 0.05

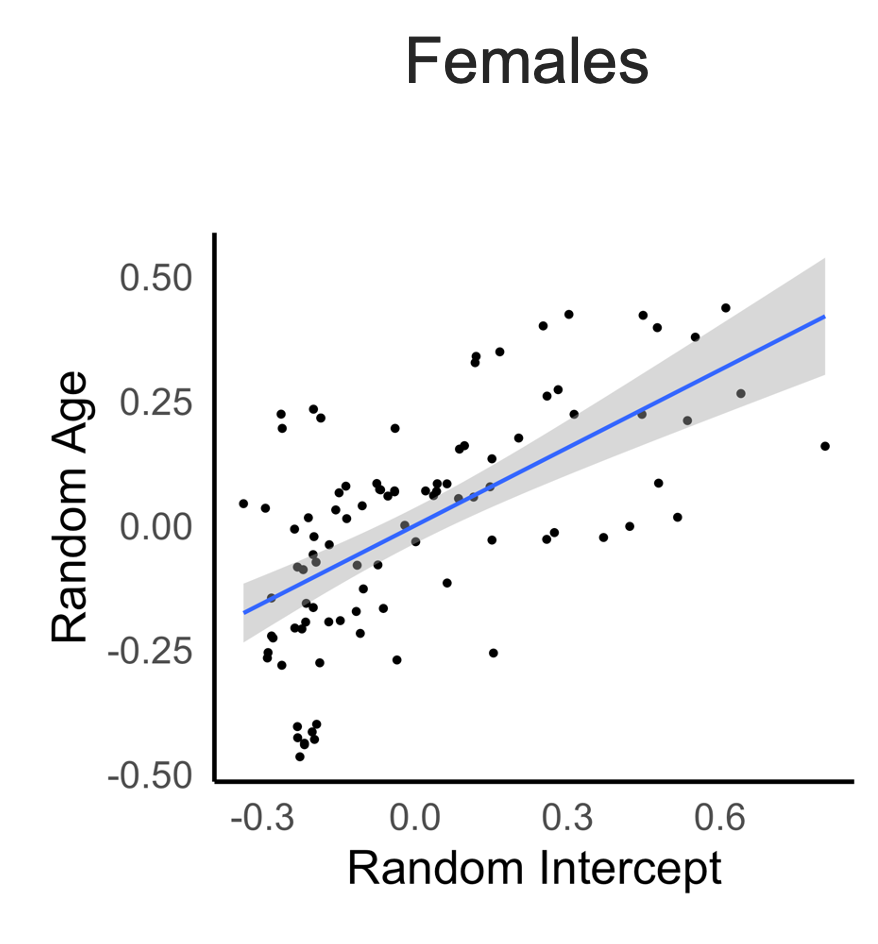

*Figure S5. Correlations between random intercept and random slopes in females*

**Age-related brain development**

As supplementary analyses, we examined the development of cortical thickness in relation to age (without consideration of pubertal stage). Specifically, we ran the following models that examined age-related development while controlling for cohort, scanner and sex:

1) thickness ~ cohort + scanner + sex + s(id, bs = “re”) + s(age, bs = “cs”, k = 3)
2) thickness ~ cohort + scanner + sex + s(id, bs = “re”) + age

The *smooth* term for age in Model 1 was corrected for multiple comparisons using False Discovery Rate 0.05, to account for the 62 regions that were examined. In addition, compareML was used to compare Models 1 and 2, to determine whether nonlinear or linear terms were a better fit to age-related trajectories.

Model comparisons revealed significant cortical thinning with age for most regions, aside from the bilateral entorhinal cortices and right medial orbitofrontal cortex. The effects were consistent when considering FDR-corrected *smooth* term for age. Model comparisons revealed linear patterns of reductions across the most cortical mantle (relative to non-linear development), characterised by an average of 1.06% change (SD: 0.38; range: -1.69 to 0.05) in cortical thickness per year (see Figure S6a). The only region exhibited a nonlinear trajectory was the right pars triangularis. Refer to Tables S8 and S9 for further detail.

As described above in the “Scanner upgrade” section, we re-ran these analyses after excluding participants above 13 years of age to deal with potential scanner confounds. As illustrated in Figure S6b, a very similar rate of change was evident across the cortex in this subsample (Mean: -1.055%, SD: 0.37, range: -1.72 to -0.01).

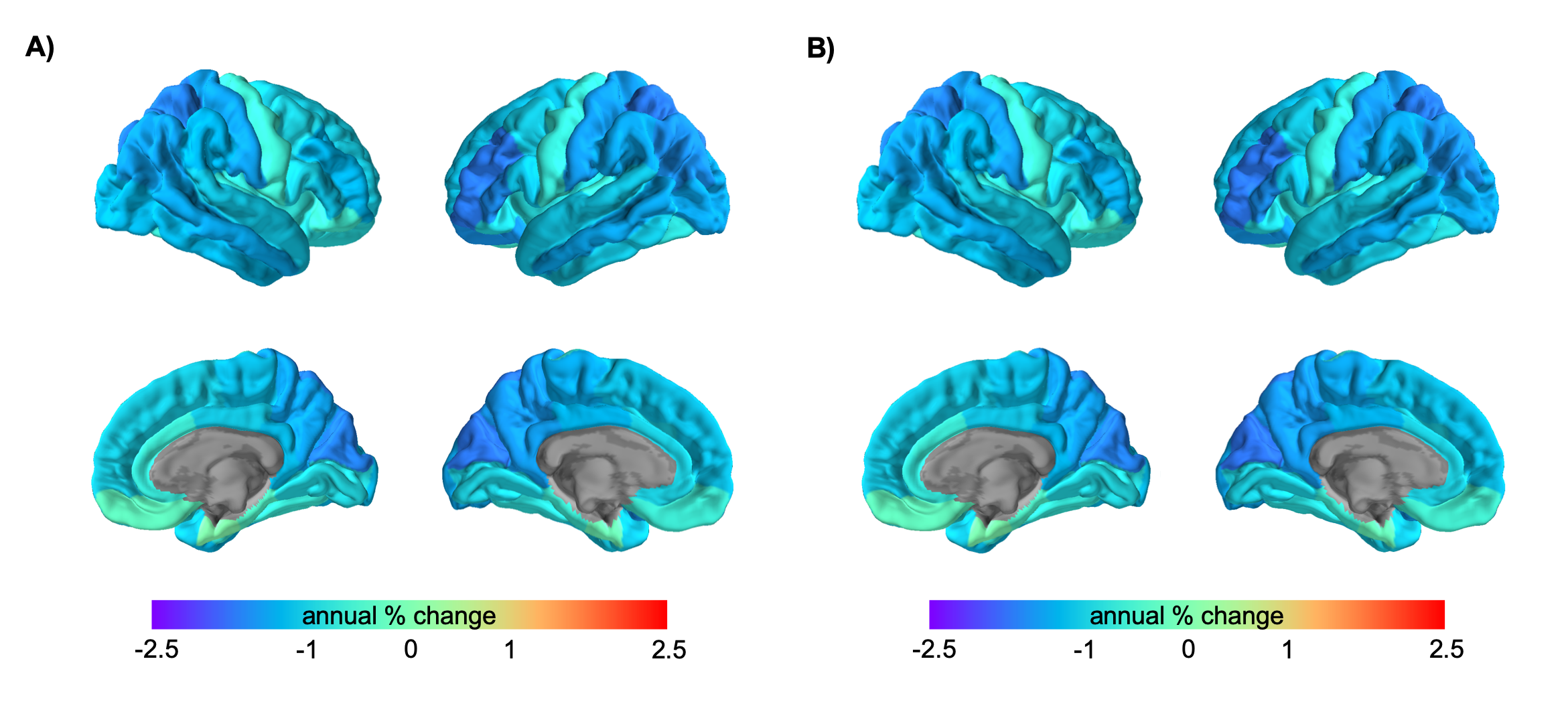

*Figure S6.* Age-related changes in cortical thickness. Values represent annualised percent change (relative to the average thickness at wave 1). A) Full sample. B) Excluding participants over 13 years of age

**Puberty-related brain development**

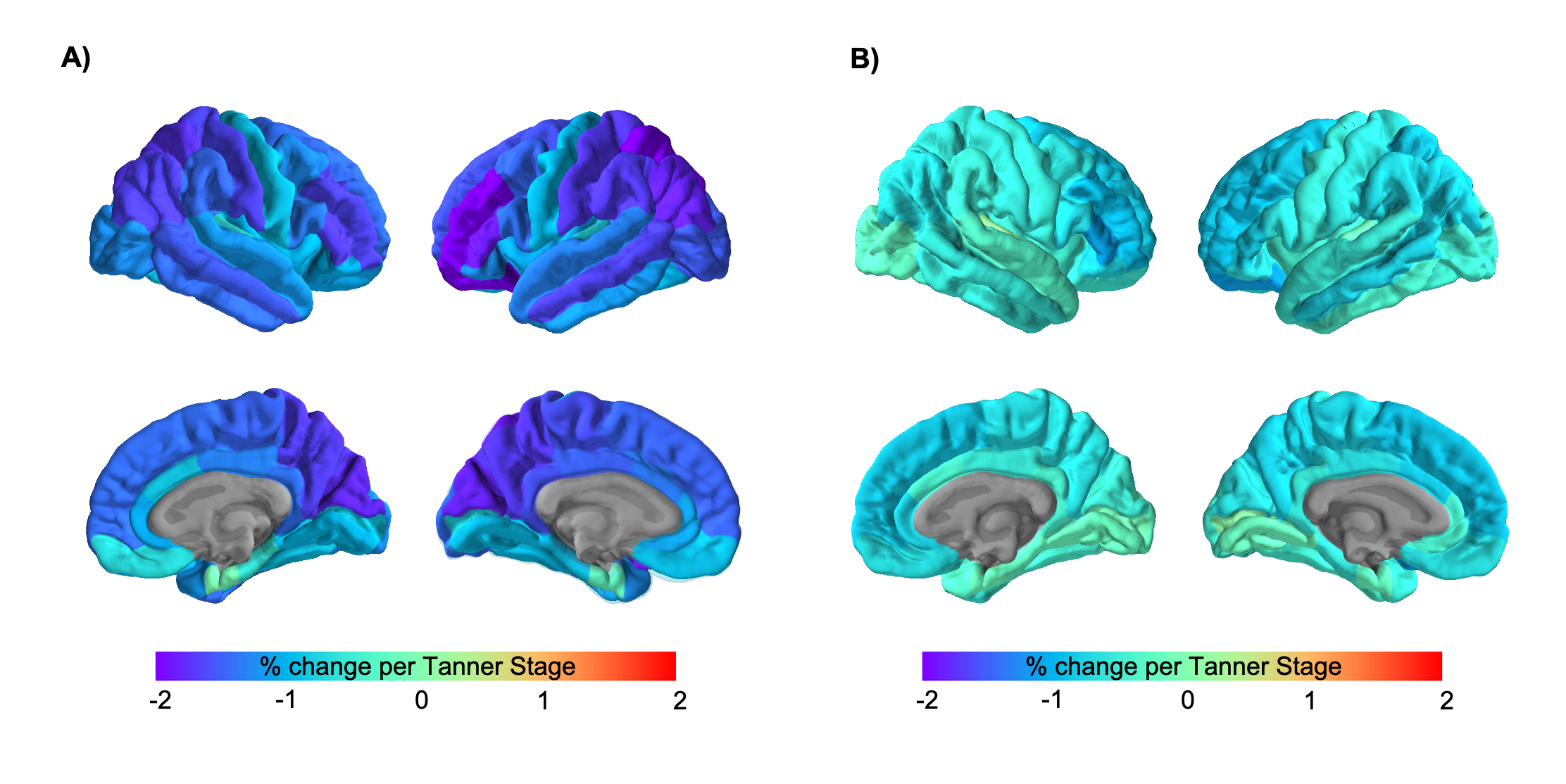

*Figure S7.* Puberty-related changes in cortical thickness, illustrated as percent change per Tanner stage, relative to the average thickness at wave 1. Comparing effect sizes without (A) and with (B) age as a covariate of non-interest.

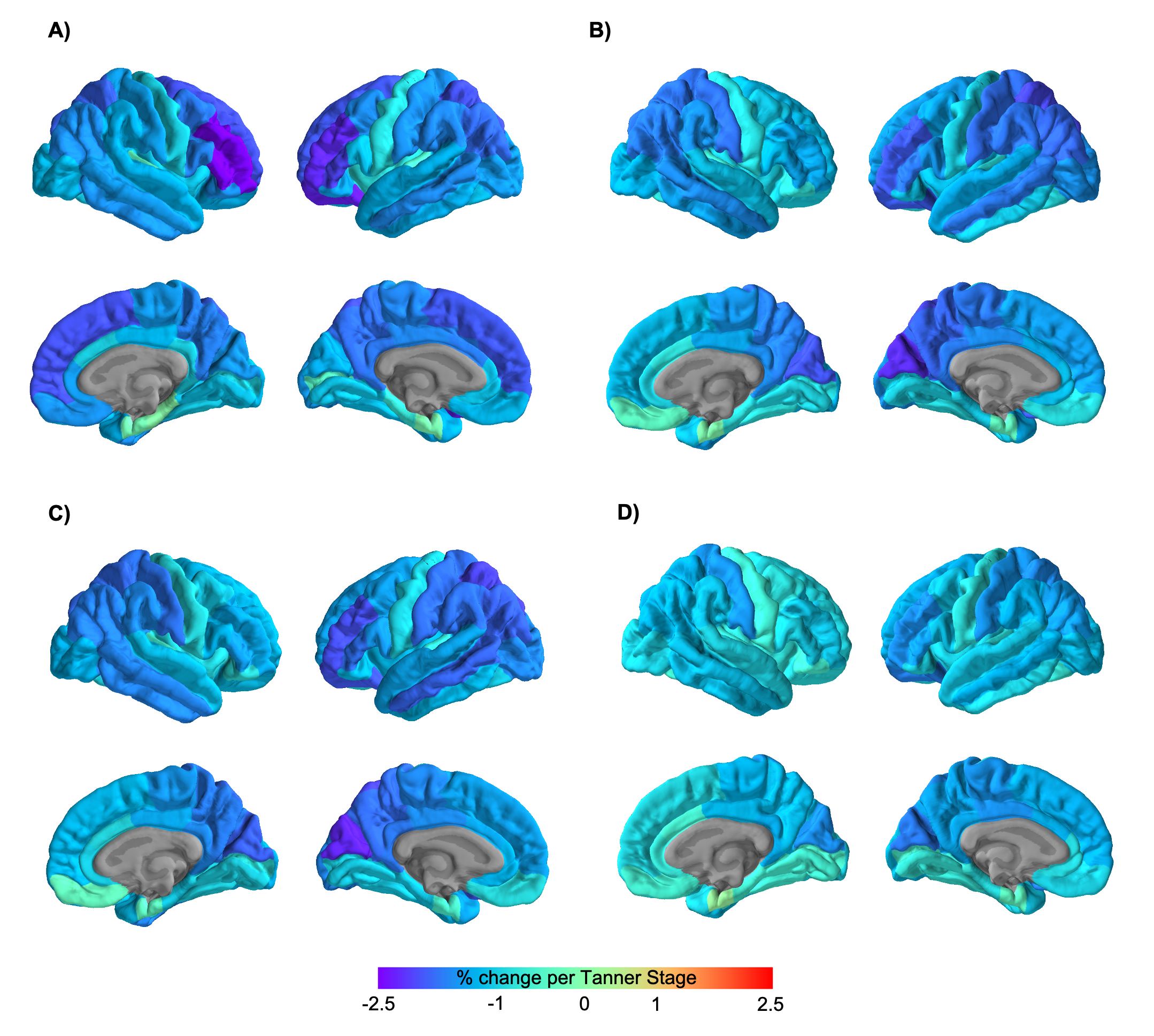
*Figure S8.* Effect sizes for puberty-related changes in cortical thickness, illustrated as percent change per Tanner stage, relative to the average thickness at wave 1. (A) Males, (B) Females, (C) Females using Tanner breast, (D) Females using Tanner hair.

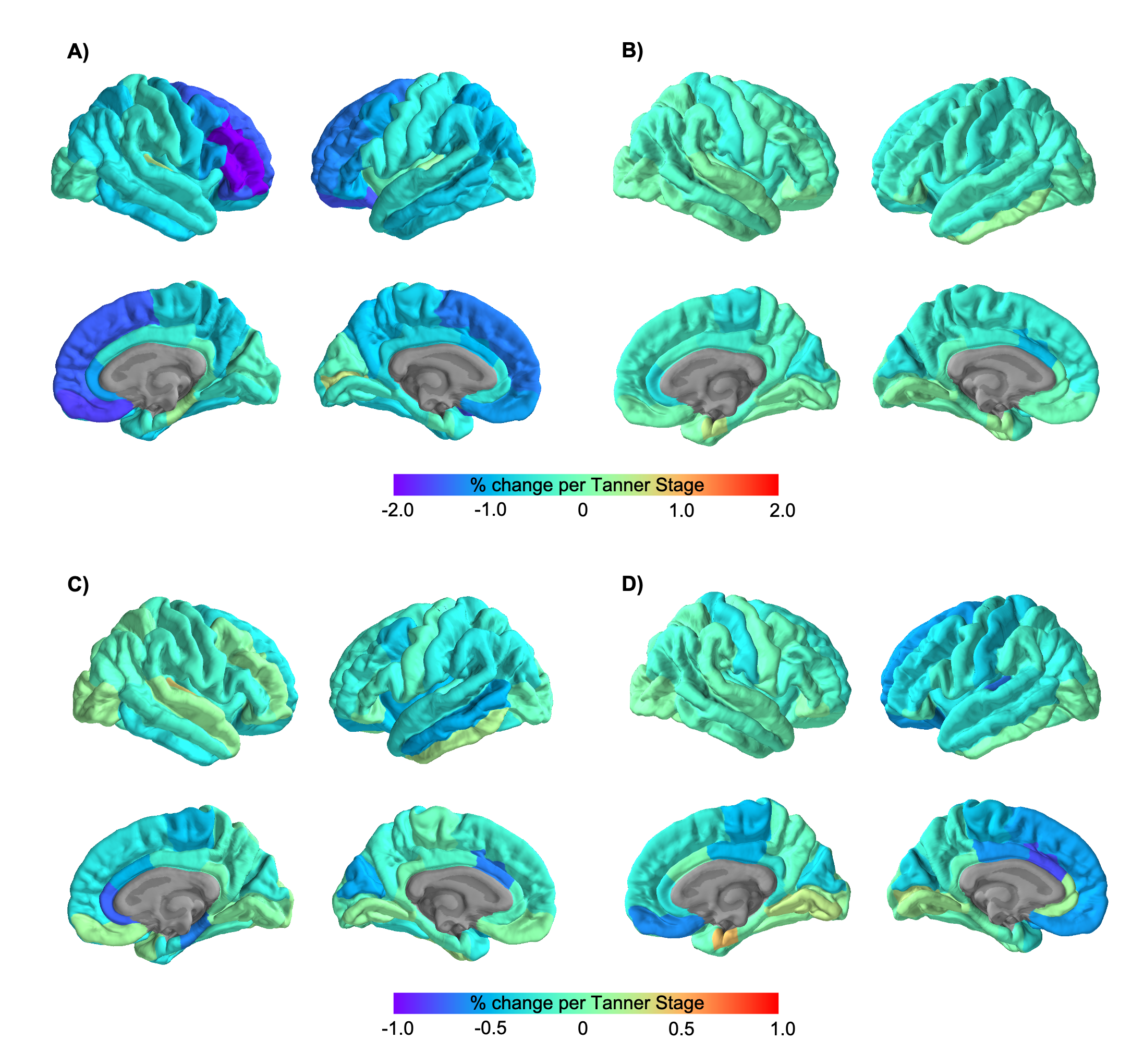
*Figure S9.* Effect sizes for puberty-related changes in cortical thickness when controlling for age, illustrated as percent change per Tanner stage, relative to the average thickness at wave 1. (A) Males, (B) Females, (C) Females using Tanner breast, (D) Females using Tanner hair.

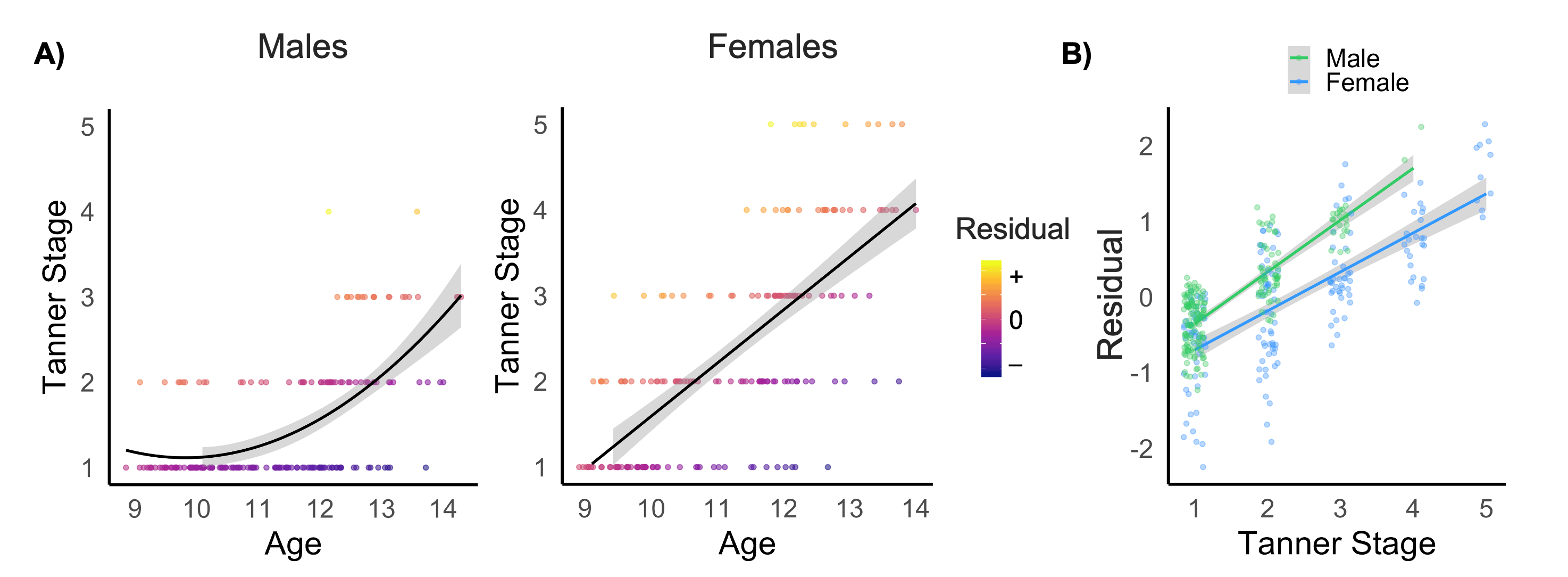

*Figure S10.* (A) The distribution of “residual puberty” (from Tanner stage ~ age) was confounded by Tanner stage, as we did not have a representation of individuals with lower residual scores at higher Tanner stages given the age span. (B) Residual puberty was positively correlated with higher Tanner stage (Males: R=0.83, p < 0.001; Females: R=0.72, p < 0.001).
